# The chemotherapeutic CX-5461 primarily targets TOP2B and exhibits selective activity in high-risk neuroblastoma

**DOI:** 10.1101/2021.02.25.432934

**Authors:** Min Pan, William C. Wright, Rich Chapple, Asif Zubair, Manbir Sandhu, Jake Batchelder, Jonathan Low, Kaley B Blankenship, Yingzhe Wang, Brittney Gordon, Payton Archer, Samuel W. Brady, Sivaraman Natarajan, Matthew J. Posgai, John Schuetz, Darcie Miller, Ravi Kalathur, Siquan Chen, Jon Patrick Connelly, M. Madan Babu, Michael A. Dyer, Shondra M. Pruett-Miller, Burgess B. Freeman, Taosheng Chen, Lucy A. Godley, Scott Blanchard, Elizabeth Stewart, John Easton, Paul Geeleher

## Abstract

Survival in high-risk pediatric neuroblastoma has remained around 50% for the last 20 years, with immunotherapies and targeted therapies having had minimal impact. Here, we identify the small molecule CX-5461 as selectively cytotoxic to high-risk neuroblastoma and synergistic with low picomolar concentrations of topoisomerase I inhibitors improving survival *in vivo* in orthotopic patient-derived xenograft neuroblastoma mouse models. CX-5461 recently progressed through phase I clinical trial as a first-in-human inhibitor of RNA-POL I. However, we also use a comprehensive panel of *in vitro* and *in vivo* assays to demonstrate that CX-5461 has been mischaracterized and that its primary target at pharmacologically relevant concentrations, is in fact topoisomerase II beta (*TOP2B*), not RNA-POL I. These findings are important because existing clinically approved chemotherapeutics have well-documented off-target interactions with TOP2B, which have previously been shown to cause both therapy-induced leukemia and cardiotoxicity—often-fatal adverse events, which can emerge several years after treatment. Thus, while we show that combination therapies involving CX-5461 have promising anti-tumor activity *in vivo* in neuroblastoma, our identification of TOP2B as the primary target of CX-5461 indicates unexpected safety concerns that should be examined in ongoing phase II clinical trials in adult patients before pursuing clinical studies in children.

## INTRODUCTION

Neuroblastoma is a pediatric cancer of the developing peripheral nervous system and the most common solid tumor in children^1^. Pediatric cancers have different mutation profiles compared to adult cancers, typically exhibiting far fewer targetable oncogene mutations^2^. Additionally, the low mutation burden and resulting lack of neoantigens means immunotherapies have had an only modest impact^3^. Consequently, the overall survival in high-risk neuroblastoma has remained around 50% for the past 20 years^4^, meaning a promising small molecule inhibitor is of tremendous interest to treat this devastating disease.

Since the release of the Cancer Cell Line Encyclopedia (CCLE)^5^ and the Genomics of Drug Sensitivity in Cancer (GDSC)^6^, large drug screening datasets in cancer cell lines have emerged as a valuable resource for identifying new therapeutic strategies to treat pediatric cancers, facilitating discoveries that could not have been made based on mutation profiles alone. For example, these datasets provided the original evidence that PARP inhibitors would be effective treating Ewing’s Sarcoma, which has since proven to have clinical activity in combination with DNA damaging agents^7, 8^. Perturbational screens in these cell lines have also motivated the development of EZH2 inhibitors in pediatric rhabdoid tumors^9^ and BRD4 inhibitors in neuroblastoma^10, 11^. However, the rapid growth of these datasets^12–14^ means systematic interrogation of the pediatric data has not been carried out, limiting the potential for prioritizing promising targets in these diseases.

Here, we mined these datasets to identify an unexpected, reproducible, selective effect of the compound CX-5461 in high-risk pediatric neuroblastoma. CX-5461 recently progressed through phase I clinical trial as a first-in-human selective inhibitor of RNA-POL I^15^, but has also been reported to target DNA G-quadruplex structures in vitro^16, 17^ and (by a single study) TOP2A^18^. Motivated by the striking drug screening results, we performed a comprehensive study in orthotopic patient-derived xenograft (PDX) bearing mice to show that CX-5461 has anti-neuroblastoma activity *in vivo* at pharmacologically relevant dosages derived using an *in vivo* pharmacokinetics modeling approach. However, we also show that CX-5461 is mischaracterized and its primary target at physiological concentrations is in fact TOP2B, which because of this gene’s unique functional role, completely changes the clinical trajectory of this compound in terms of patient stratification, drug combinations, and likely side-effect profiles. Critically, *TOP2B* is highly expressed in some normal cells and off-target drug interactions with this gene have previously been implicated in late-emerging therapy-induced acute leukemias^19, 20^ and cardiotoxicity^21^, which often lead to death. Thus, while CX-5461 has promising anti-tumor activity *in vivo*, our results indicate that it also has the potential to cause previously unanticipated patient harm, which should be investigated in adults before clinical studies are proposed in children.

## RESULTS

### CX-5461 exhibits selective and potent activity against neuroblastoma cell lines

We first devised a quantitative metric to identify compounds with selective activity against neuroblastoma cell lines in large drug screening datasets (similar to Durbin *et al.*^10^). For each drug, we ranked cell lines by their IC_50_, identified the rank of the median neuroblastoma cell line, then normalized this value to a 0-1 scale by dividing by the total number of cell lines screened against this drug—we refer to this as a “Selectivity Score”. We calculated Selectivity Scores for each drug in the most recent release of the Sanger Institute’s GDSC cell line drug screening dataset, where a total of 265 drugs were screened against 1,001 cancer cell lines^22^, including 31 neuroblastoma cell lines, the most of currently available datasets. Drugs with the highest Selectivity Score included several already in investigation in neuroblastoma (Fig. 1A; Table S1), for example, ranked #3 and #4 were IGF1R inhibitors (linsitinib and BMS-754807)^23^.

**Figure 1.**
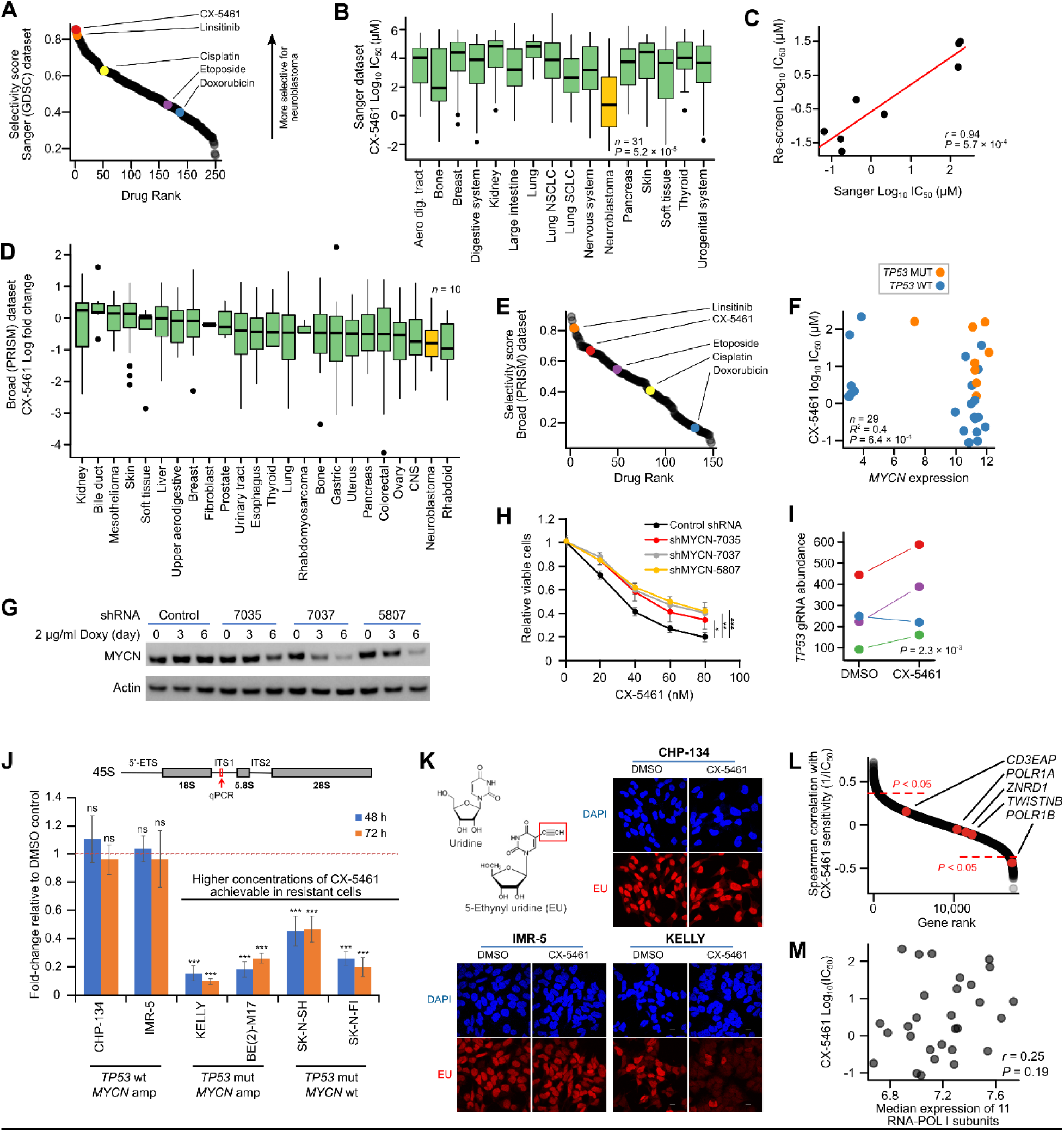
Neuroblastoma cell lines are selectively sensitive to CX-5461. A. Waterfall plot showing summary score representing selectivity for neuroblastoma cell lines plotted for each of 265 drugs screened by GDSC, where the y-axis is the observed scores, and the x-axis is the drug rank. A higher value on the y-axis implies greater selective activity of the drug against neuroblastoma cell lines. CX-5461 is highlighted in red. B. Boxplot of log_10_(IC_50_) values for CX-5461 for all non-hematological cancer cell lines screened by GDSC (see Fig. S1A). C. Scatterplot of log_10_(IC_50_) values for CX-5461 for 8 cell lines screened by GDSC (x-axis) and re-screened i this study (y-axis). D. Boxplot of log_10_(fold change) values for CX-5461 for all cell lines screened by PRISM. Fold-changes represent the difference in cell viability of drug treated vs. control cells in the PRISM assay, estimated by sequencing each cells unique barcode. A lower fold-change implies higher drug effectiveness. Note: there were no rhabdoid cell lines in the GDSC discovery dataset. E. Waterfall plot showing summary score representing selectivity for neuroblastoma cell lines plotted for each of 148 drugs screened in PRISM (shown for drugs screened by both PRISM and GDSC), where the y-axis are the observed scores, and the x-axis is the drug rank. F. Scatterplot showing *MYCN* expression level (x-axis) against log_10_(IC_50_) values for CX-5461 in GDSC (y-axis). The points are colored by *TP53* mutation status. G. Western blot showing MYCN protein levels following *MYCN* knockdown using 3 independent shRNAs in CHP-134 cells. β-Actin was used as loading control. Doxy, doxycycline. H. Viability of CHP-134 cells following *MYCN* knockdown under treatment with CX-5461. Cells were transduced with one of three independent *MYCN* shRNAs or a negative control shRNA. After 6 days of incubation in medium with 2 µg/ml doxycycline, cells were treated with CX-5461 for 3 days. Cell viability was measured with MTS. Data represent mean ± SD of 3 independent experiments. \**P* < 0.05, \*\**P* < 0.01, \*\*\**P* < 0.001. I. Stripchart showing the relative abundance of 4 independent *TP53* guide RNAs in a genome-wide CRISPR screen (y-axis) in either a DMSO or CX-5461 treated CHP-134 neuroblastoma cell line. J. Pre-rRNA 45S expression (y-axis) in CX-5461 treated cell lines relative to DMSO, determined by RT-qPCR with primers located in an internal transcribed spacer (ITS) region of the rRNA transcript. Data represent mean ± SD of 3 independent experiments. \*\*\**P* < 0.001; ns, no significant difference to DMSO control. CX-5461 concentration: CHP-134, 0.2 µM; IMR-5, 0.05 µM; KELLY, 2 µM; BE(2)-M17, 10 µM; SK-NSH, 2 µM; SK-N-FI, 20 µM. K. EU incorporation assay to assess global nascent RNA transcription. CHP-134, IMR-5 and KELLY cells were treated with CX-5461 for 24 h. 1 mM EU was added at 30 min (CHP-134, IMR-5) or 1 h (KELLY) before cell fixation. Nascent RNA was labeled with EU (red). Cell nuclei were stained with DAPI (blue). CX-5461 concentration: CHP-134, 0.2 µM; IMR-5 0.05 µM; KELLY, 2 µM. Scale bar = 10 μm. L. Waterfall plot showing Spearman’s correlation of the expression of all genes in GDSC with the reciprocal of CX-5461 IC_50_ (y-axis) in 29 neuroblastoma cell lines. Higher values on the y-axis imply that high expression of a gene is associated with greater sensitivity to CX-5461. Genes unique to the RNA-POL I complex (those not shared with RNA-POL II) are highlighted in red. M. Scatterplot showing the correlation of the median expression level of 11 genes of the RNA-POL I complex (x-axis) for which a gene expression estimate was available in GDSC with CX-5461 log_10_(IC_50_) (y-axis) in 29 neuroblastoma cell lines.

Surprisingly, the second most selective drug for neuroblastoma was CX-5461, a compound that recently completed phase I clinical trial as a first-in-human selective inhibitor of RNA-POL I, but is not currently in clinical investigation in neuroblastoma^4^. The selective activity was striking (Fig. 1B, Fig. S1A, *P* = 5.2 × 10^-6^ from t-test of neuroblastoma IC_50_ values vs all other cell lines) with almost half of neuroblastoma cell lines achieving an IC_50_ in the nanomolar range. We obtained and rescreened 8 of the GDSC cell lines with CX-5461 and achieved highly consistent results (Fig. 1C, *r* = 0.94, *P* = 5.7 × 10^-4^; Fig. S1B-C). Additionally, CX-5461 was included in PRISM, a recent independent study^12^ that screened 4,518 drugs against 578 cancer cell lines using a new high-throughput multiplex sequencing-based assay. Although only 10 neuroblastoma cell lines were included, neuroblastoma remained the second most sensitive group to CX-5461 among the 24 cancer types screened (Fig. 1D). The overall trend for Selectivity Scores was also recapitulated in PRISM (Fig. 1E; Fig. S1D), with CX-5461 remaining among the most selective drugs for neuroblastoma cell lines.

To deduce whether CX-5461 may favor specific subclasses of neuroblastoma, we next sought to identify genetic predictors of CX-5461 response in these neuroblastoma cell lines. Using the GDSC’s exome sequencing data we identified *MYCN* amplification and *TP53* mutation as independently predictive of response (*P* = 6.4 × 10^-4^ from a multivariate linear model; *R^2^* = 0.4). The effect sizes were large; *TP53* mutation was associated with a 17.45 ± 4.7 (mean effect +/- 95% CI) fold increase in IC_50_, while *MYCN* amplification was associated with a 10.9 ± 5.6-fold decrease in IC_50_. We tested these predictions *in vitro* using short hairpin RNA (shRNA) knockdown of *MYCN* (*P* < 0.05 for each of 3 independent shRNAs, Fig. 1G-H) and CRISPR knockout of *TP53* (*P* = 2.3 × 10^-3^, Fig. 1I) in CHP-134 (*MYCN* amp, *TP53* wt.) neuroblastoma cells, identifying a causal relationship with CX-5461 IC_50_ in both cases. We recalculated the Selectivity Scores for only *MYCN* amplified *TP53* wild-type neuroblastoma cell lines— representative of high-risk disease—where CX-5461 was the #1 most selective compound for these cell lines (Fig. S1E & S1F). Thus, CX-5461 is of particular interest in the high-risk subgroup of neuroblastoma, where *MYCN* amplification is the primary driver and patient outcomes are extremely poor.

### CX-5461 kills neuroblastoma cells at low concentrations that do not inhibit RNA-POL I

CX-5461 recently completed phase I clinical trial as a selective inhibitor of RNA-POL I^24^. To confirm that this was indeed the mechanism-of-action against neuroblastoma cells, we performed a standard assay to determine whether CX-5461 interferes with RNA-POL I activity^25^—using qPCR to the short-lived internal transcribed spacer (ITS) region of the 45S rRNA transcript. We applied this in CX-5461-treated CHP-134 and IMR-5 (both highly CX-5461 sensitive) neuroblastoma cell lines. Surprisingly, we observed no inhibition of RNA-POL I activity in these cell lines at the low sub-micromolar concentrations at which CX-5461 was effective (Fig. 1J, S1G-H). However, in four CX-5461-resistant neuroblastoma cell lines, where higher CX-5461 concentrations could be tolerated, we observed inhibition of RNA-POL I (Fig. 1J). This is consistent with RNA-POL I inhibition emerging either as an off-target effect or as a consequence of inhibiting the primary drug target. We validated these observations using a 5-ethynyl uridine (EU) incorporation assay, where again there was no inhibition of RNA synthesis at the low concentrations of CX-5461 that were effective against CHP-134 and IMR-5 cells (Fig. 1K). The resistant cell line KELLY, which could be treated with higher concentrations of CX-5461, again showed clear inhibition of RNA-POL I. We also identified no significant associations between the baseline expression levels of RNA-POL I complex genes and CX-5461 IC_50_ in GDSC neuroblastoma cell lines (Fig. 1L) or between the median expression of RNA-POL I complex genes and response to CX-5461 (Fig. 1M). Finally, treatment with CX-5461 for 48 or 72 hours did not decrease the levels of mature cellular rRNA in neuroblastoma cells (Fig. S1I-J). These results indicate that CX-5461’s anti-cancer activity may not exclusively result from selective inhibition of RNA-POL I. Instead, these findings suggest that CX-5461 may be active against neuroblastoma cell lines by one or more additional mechanisms independent of impacts on rRNA synthesis.

### CX-5461’s is a topoisomerase II inhibitor

The genome-wide CRISPR knockout screen has emerged as a powerful, unbiased assay for determining the mechanism-of-action of small molecules^26^. The workflow involves culturing cells stably expressing Cas9 with or without the drug of interest, performing genome-wide CRISPR gene knockout, and comparing the relative abundance of different guide RNAs in drug-treated vs control cells; this acts as a proxy for the proportion of surviving cells that have undergone each gene knockout (Fig. 2A). We performed such a screen in CHP-134 neuroblastoma cells treated with 0.04 μM CX-5461 or DMSO control, 205 allowing us to identify genes required for growth of CHP-134 cells in the presence of CX-5461. Notably, none of the top hits had any obvious relationship to RNA-POL I or DNA G-quadruplexes, however, there was a clear enrichment of genes related to DNA-damage response (Fig. 2B, Fig. S2A-B). Remarkably, all three of the most significant knockouts had a direct relationship to topoisomerase II (Fig. 2C-E): TDP2 catalyzes the removal of the covalent bonds between topoisomerase II and DNA^27^, LIG4 is a DNA ligase that rejoins double-strand breaks caused during normal topoisomerase II activity^28^, and inhibition of FZR1 has previously been shown to sensitize cells to topoisomerase II inhibitors^29^. We queried a recently published resource^26^ that performed CRISPR knockout screens in human retinal epithelial cells treated with 31 DNA damaging agents and confirmed that *TDP2*, *LIG4*, and *FZR1* knockouts all sensitize these cells to the known topoisomerase II inhibitors doxorubicin and etoposide (Fig. 2F-G). These results suggest that CX-5461’s primary target at pharmacologically relevant concentrations is in fact topoisomerase II. This is especially plausible given that CX-5461 is a fluoroquinolone derivative^30^ and maintains structural similarity to these antibiotics, which target bacterial DNA gyrase—an ortholog of human topoisomerase II^31^.

**Figure 2.**
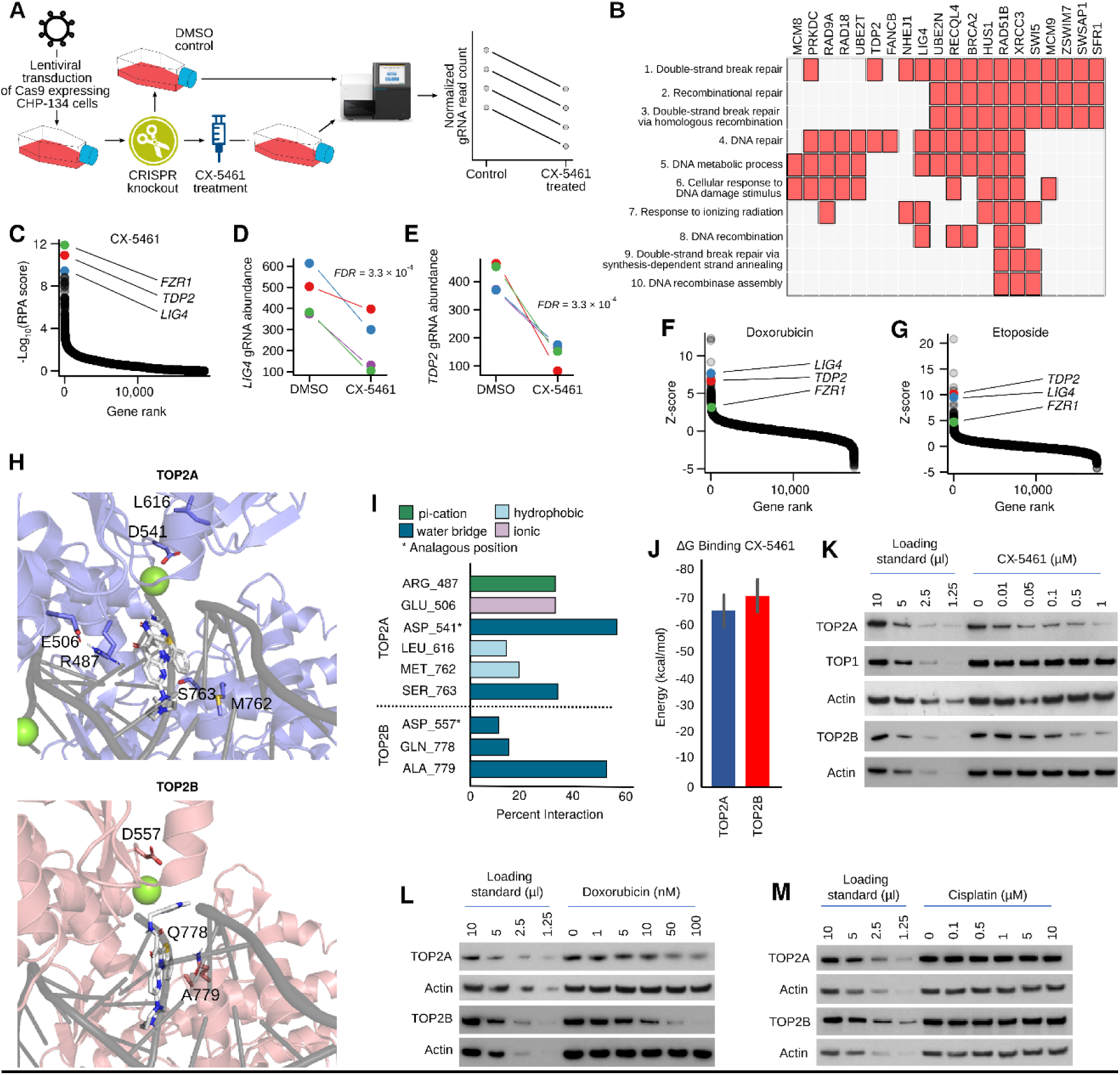
CX-5461 is a topoisomerase II inhibitor. A. Schematic of CRISPR knockout screening strategy. CX-5461 (0.04µM) and DMSO (no treatment negative control) treated CHP-134 cells were subjected to whole-genome CRISPR screen. B. Clustergram generated by Enrichr showing the assignment of statistically significant (red boxes) gen knockouts to the most highly enriched GO (2018) Biological Processes (y-axis). Genes with an FDR < 0.05 (n = 40) were used as input. C. Waterfall plot highlighting the top hits in our genome-wide CRISPR knockout screen in CHP-134 cells. Genes are ranked (x-axis) by −log_10_(RPA score) (y-axis). D. Stripchart showing the relative abundance of 4 independent *LIG4* guide RNAs in our genome-wide CRISPR screen (y-axis) in either DMSO or CX-5461 treated CHP-134 cells. E. Like (D) but for *TDP2*. F. Waterfall plot summarizing CRISPR knockout screening data obtained from Oliveri *et al.* for the topoisomerase II inhibitor doxorubicin. Higher values on the y-axes imply that knockout has sensitized cells to doxorubicin. *TDP2, LIG4,* and *FZR1* are highlighted in red, blue and green respectively. G. Like (F) but results shown for the topoisomerase II inhibitor etoposide. H. Representative poses from equilibrated molecular dynamics simulations of CX-5461 bound to TOP2A (top) or TOP2B (bottom). DNA strands are shown in gray. CX-5461 is displayed in white, and its heteroatoms colored blue (nitrogen), red (oxygen), and yellow (sulfur). Green represents magnesium ions. I. Percentage of CX-5461 interactions with TOP2A or TOP2B simulations. Residues interacting with either protein are listed if they conferred ≥ 10% interaction with CX-5461. Bar colors correspond to type of interaction. (*) indicates residues in the same position when the two proteins are superimposed. J. Weighted average of the relative binding affinity for CX-5461 is plotted for TOP2A (65.2 kcal/mol) and TOP2B (−70.7 kcal/mol). Error bars represent the standard deviation of the weighted averages (*P* = 0.01 from t-test). K-M. Band-depletion assay for CX-5461 (K), doxorubicin (L), and cisplatin (M) treated CHP-134 cells. Cells were exposed to agents at their approximate IC_50_ for 4 h. Cell lysates were collected for SDS-PAGE. Cell lysates from untreated cells were used as loading standard.

To investigate further whether CX-5461’s structure is consistent with that of a topoisomerase II inhibitor, we performed molecular dynamics simulations, using existing TOP2A and TOP2B protein structures. The *in silico* binding of CX-5461 was consistent with that of a classical topoisomerase II poison: This is the most common mechanism for topoisomerase II inhibitors (including fluoroquinolones, etoposide, and doxorubicin), where the drug traps the topoisomerase protein covalently bound to DNA, leading to DNA damage and cell death^32^. CX-5461 interacts with the DNA breakpoint, collectively surrounded by the TOP2A/B DNA binding domains (Fig. 2H). We also found that CX-5461 interactions differ between TOP2A and TOP2B, interacting with largely non-analogous amino acids of the respective proteins (Fig. 2I). We calculated the relative binding affinity of the CX-5461 ligand to the TOP2A/B proteins and noted a slightly increased affinity of the ligand to TOP2B. (Fig. 2J).

We performed several orthogonal assays to test the hypothesis that CX-5461 is a topoisomerase II poison. We first performed a band depletion assay^33^, to assess the stabilization of TOP2-DNA covalent complexes (called the TOP2 cleavable complex (TOP2cc)), following CX-5461 treatment. This assay showed a clear depletion of free TOP2A and TOP2B protein, consistent with these proteins being trapped to DNA following treatment with CX-5461 (Fig. 2K). The decrease of free TOP2 was dose-dependent and comparable to the positive control TOP2 poison doxorubicin (Fig. 2L). The cisplatin-treated negative control showed no change (Fig. 2M). CX-5461 did not affect TOP1 (Fig. 2K), suggesting its activity is selective to TOP2. We next used a complementary assay, the rapid approach to DNA adduct recovery (RADAR) assay, to further assess the ability of CX-5461 to form TOP2cc^34^. This revealed CX-5461 treatment-induced TOP2cc formation, for both TOP2A and TOP2B, in all four neuroblastoma cell lines tested (Fig. S2C), again consistent with CX-5461 trapping TOP2 to DNA and behaving as a classical topoisomerase II poison.

### Low sub-micromolar concentrations of CX-5461 induce cell cycle arrest and kill neuroblastoma cells by causing DNA damage and cell death by apoptosis

The mechanism of drug-induced cell death by classical topoisomerase II poisons, such as etoposide and doxorubicin is well understood. These drugs trap topoisomerase protein to DNA, and these DNA-topoisomerase complexes are converted to DNA double-strand breaks when encountered by the DNA replication or transcription machinery, leading to cell death by apoptosis^35^.

To investigate whether this was the mechanism of cell death in neuroblastoma cells susceptible to sub-micromolar concentrations of CX-5461, we first used flow cytometry to quantify neuroblastoma cells staining positive for annexin V and propidium iodide (PI), canonical markers of apoptosis and cell death respectively, following CX-5461 treatment at 0.1 μM, 0.2μM and 0.5 μM over 48h. These assays showed that CX-5461 was causing cell death by apoptosis at these low concentrations in CX-5461 sensitive CHP-134 and IMR-5 neuroblastoma cells (Fig. 3A-3C). Induction of cell death by apoptosis at low concentrations of CX-5461 was confirmed by Western blot for cleaved PARP (c-PARP, Fig. 3D) in CHP-134 and IMR-5 cell lines.

**Figure 3.**
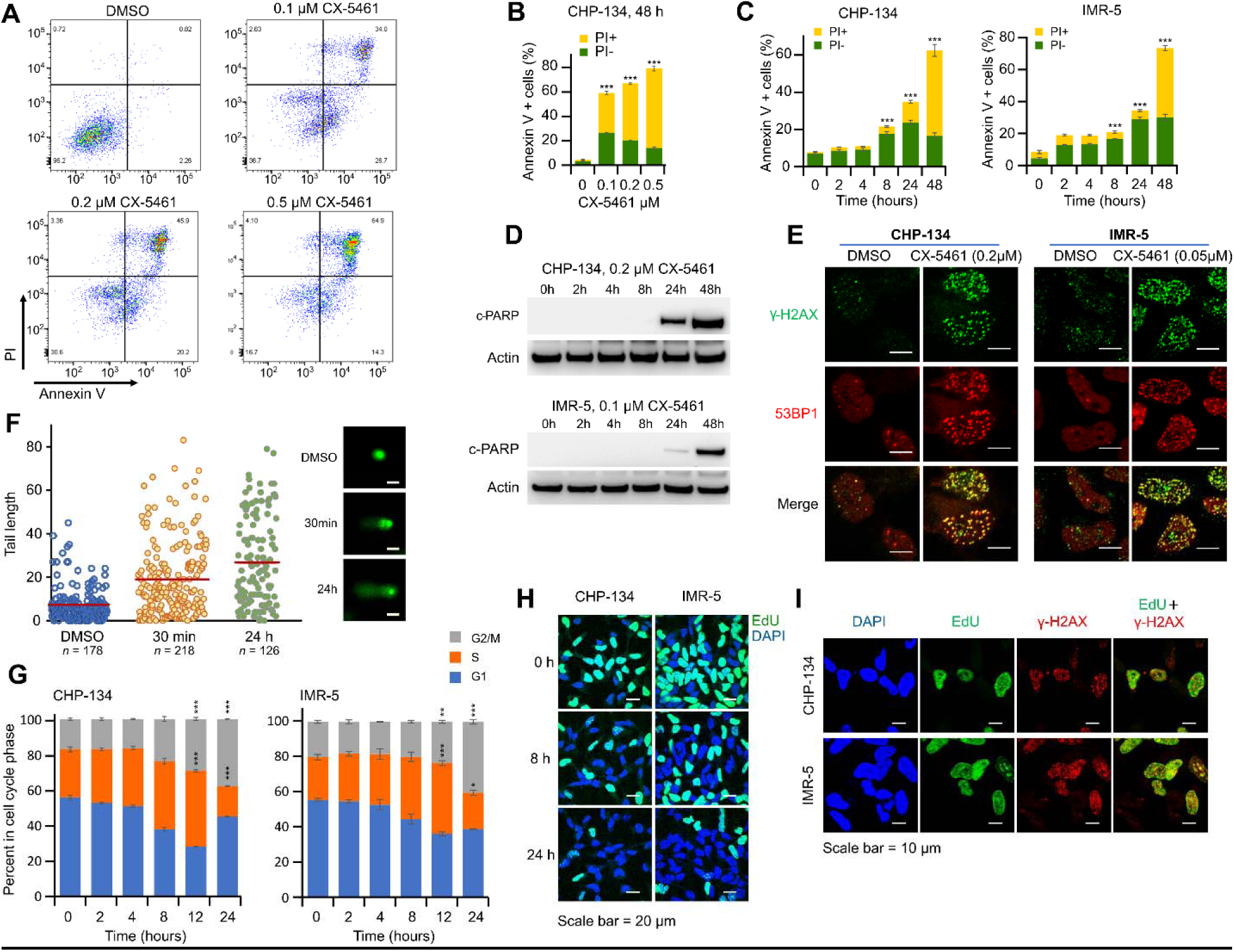
CX-5461 causes DNA damage and cell death by apoptosis in neuroblastoma cells at low concentrations that do not inhibit RNA POL I. A. Flow cytometry data showing apoptosis in CHP-134 cells. The cells were treated with CX-5461 for 48 with indicated concentrations. Induction of cell apoptosis was measured by annexin V (x-axis) and cell death by propidium iodide (PI) staining (y-axis). B. Quantification of data in Fig. 2E. The y-axis shows the percentage of annexin V positive cells, with the proportion of PI positive cells shown in yellow, for increasing concentrations of CX-5461 (x-axis). Data represent the mean ± SD of 3 independent experiments. \*\*\**P* < 0.001. C. Quantification of flow cytometry data showing apoptosis up to 48 h in CHP-134 (left panel) and IMR-5 (right panel) cells. CHP-134 cells were treated with 0.2 µM and IMR-5 cells with 0.1 µM of CX-5461. The y-axis shows the percentage of annexin V positive cells, with the proportion of PI positive cells shown i yellow (x-axis). Data represent the mean ± SD of 3 independent experiments. \*\*\**P* < 0.001. D. Western blots for cleaved PARP (c-PARP). Cells were treated with CX-5461 for the time an concentration indicated. β-Actin was used as a loading control. E. Co-localization of 53BP1 and γ-H2AX. Cells were treated with DMSO or CX-5461 (0.2 µM for CHP-13 and 0.05 µM for IMR-5) for 24 h, cells were stained with 53BP1 (red) and γ-H2AX (green) antibodies for immunofluorescence. Scale bar = 10 μm. F. Comet assay in CHP-134 cells. Cells were treated with DMSO or 0.2 µM CX-5461 for 30 min or 24 h (x-axis) before electrophoresis. Tail length (left) and representative images (right) are shown. The red line indicates the mean value of tail length in each group. Scale bar = 5 µm. G. Bar graph showing results of cell cycle analysis. CHP-134 and IMR-5 cells were treated with CX-5461 (CHP-134, 0.2 µM; IMR-5, 0.05 µM) for the time indicated (x-axis) and stained with PI for flow cytometry analysis. The proportion of cells in each cell cycle phase (y-axis) was determined by DNA content. Data represent the mean ± SD of 3 independent experiments. \**P* < 0.05, \*\**P* < 0.01, \*\*\**P* < 0.001. H. CHP-134 and IMR-5 cells were treated with 0.2 µM CX-5461 and 0.05 µM CX-5461 respectively for the time indicated. 2 hours before cell fixation, 10 µM EdU was added to the medium. Blue, DAPI; green, EdU. Scale bar = 20 µm. I. CX-5461 induced DNA damage (γ-H2AX, red) was enriched in cells with active DNA synthesis (positive for EdU (green) labeling). CHP-134 cells were treated with 20 μM EdU and 0.2 μM CX-5461 for 3 h. IMR-5 cells were treated with 10 µM EdU and 0.1 µM CX-5461 for 3 h. Immunofluorescence was then carried out in these cells. Scale bar = 10 μm.

We next investigated CX-5461’s DNA damaging effects, using immunofluorescence for canonical DNA damage markers γ-H2AX, 53BP1 (Fig. 3E, Fig. S3A), and RPA (Fig. S3B) in CHP-134 and IMR-5 cell lines. Indeed, these markers were evident following treatment with nanomolar concentrations of CX-5461 for 24 hours. We confirmed DNA damage using a single-cell alkali gel electrophoresis (comet) assay in CHP-134 cells treated with 0.2 µM for 30 minutes or 24 hours, with DNA damage evident at 30 minutes and increasing at 24 hours (Fig. 3F, Fig. S3C). Furthermore, using cell cycle analysis and an EdU incorporation assay, we found evidence of S phase cell cycle arrest (Fig. 3G) and deceleration of active DNA synthesis (Fig. 3H), started 8 h post CX-5461 treatment. Moreover, γ-H2AX staining was heavily enriched in cells with active DNA synthesis (Fig. 3I). Together, these data suggest that DNA replication stress was induced and DNA damage was generated during DNA replication. Western blot revealed phosphorylation of cell cycle checkpoints CHK1, CHK2, and p53(S15) in CX-5461 treated CHP-134 and IMR-5 cells, indicating activation of the DNA-damage response (Fig. S3D-EI). We also found that cells surviving over a 24 h treatment with CX-5461 accumulated in the G2/M phase of cell cycle (Fig. 3G), consistent with arrested cells undergoing DNA repair. In summary, the DNA damaging effects of low sub-nanomolar concentrations of CX-5461 (not sufficient to inhibit RNA-POL I) follow the hallmark DNA-damaging activities of topoisomerase II poisons.

### CX-5461 primarily targets the TOP2B paralog

The findings above initially seemed to be consistent with single isolated study, which used a computational systems biology approach to predict TOP2A as CX-5461’s primary target^18^. This is exciting as TOP2A is one of the most successful drug targets in cancer^36^; however, TOP2A as CX-5461’s primary target leaves critical inconsistencies. Why did other topoisomerase II poisons (etoposide, doxorubicin) not also show selective activity against neuroblastoma cell lines (Fig. 1A & 1E)? Moreover, why has CX-5461 been shown to inhibit RNA-POL I in many different cell types, including non-cycling cells^37^, which do not express TOP2A^38^? As was hinted at in our molecular dynamic simulations, the answers stem from CX-5461’s interaction with TOP2B—which despite structural similarity is highly functionally distinct from *TOP2A*^19^, thus completely altering the clinical trajectory of this compound.

We first observed that although *TOP2A* and *TOP2B* were both overexpressed in neuroblastoma cell lines (Figure S4A-B), the magnitude of *TOP2B* overexpression was substantially larger. Whereas the median expression of *TOP2A* was 31.3% higher than the median in all other cancer cell lines in DepMap RNA-seq data (*P* = 2 × 10^-3^, Figure 4A), the median expression of *TOP2B* was 87.1% higher (*P* = 1.3 × 10^-7^, Figure 4B). Additionally, *TOP2B* expression was among the most highly negatively correlated genes with CX-5461 IC_50_ in GDSC neuroblastoma cell lines (*r* = -.58, *P* = 1 × 10^-3^, Fig. 4C), and this was the 2^nd^ highest negative correlation with *TOP2B* of all drugs screened (Fig. 4D). Based on these observations, we hypothesized that CX-5461’s cytotoxicity may occur through preferential targeting of TOP2B, unlike existing TOP2 poisons etoposide and doxorubicin, whose effects primarily result from trapping TOP2A on DNA.

**Figure 4.**
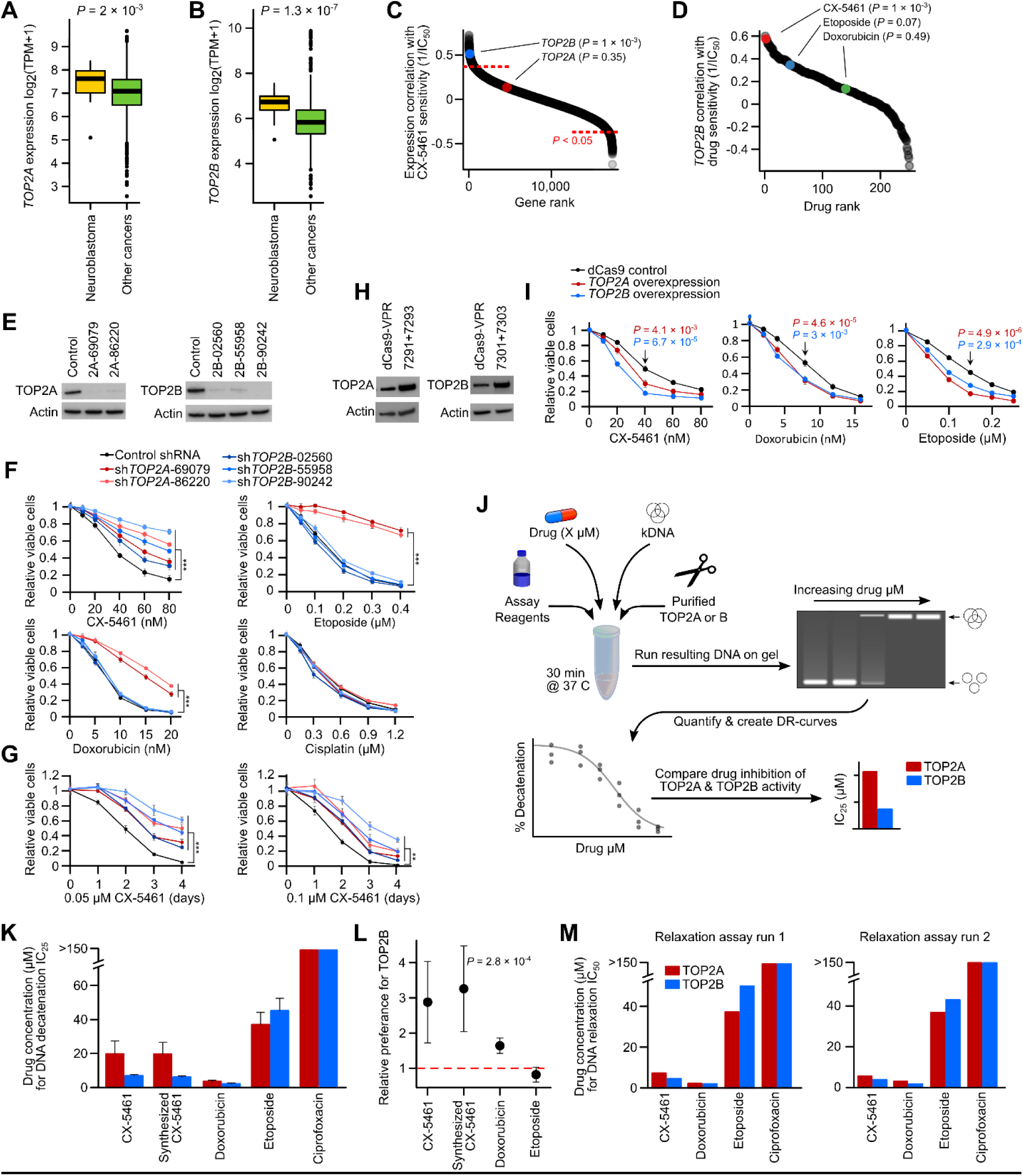
CX-5461 primarily targets the TOP2B paralog. A. Boxplot showing the log_2_(TPM+1) expression of *TOP2A* (y-axis) in DepMap neuroblastoma cell lines compared to all other DepMap cell lines, measured by RNA-seq. B. Like (A) but for *TOP2B*. C. Waterfall plot showing Spearman correlations of the expression of all genes with CX-5461 sensitivity in GDSC neuroblastoma cell lines, with *TOP2A* and *TOP2B* highlighted in red and blue respectively. D. Waterfall plot for the Spearman correlation of *TOP2B* expression with sensitivity (1/IC_50_) of all drugs screened in GDSC neuroblastoma cell lines. CX-5461, etoposide and doxorubicin are highlighted in red, blue and green respectively. E. Western blots for *TOP2A* (left) and *TOP2B* (right) following shRNA knockdown. CHP-134 cells were transduced with *TOP2A* or *TOP2B* shRNAs (2 shRNAs for *TOP2A* and 3 shRNAs for *TOP2B*). A control shRNA was used with scramble target in human cells. Cells were cultured in medium with 0.5 µg/ml puromycin and 2 µg/ml doxycycline for 3 days, then cell lysates were collected for western blotting. β-Actin was used as a loading control. F. *TOP2A* (red) and *TOP2B* (blue) knockdown CHP-134 cells and negative control (black) cells were cultured in medium with 2 µg/ml doxycycline for 3 days, then, the indicated concentration of CX-5461, etoposide, doxorubicin or cisplatin was added to the medium for 72 hours. Cell viability was determined by MTS assay. Data represent mean ± SD of 3 independent experiments. \*\*\**P* < 0.001. G. *TOP2A* (red) and *TOP2B* (blue) knockdown CHP-134 cells and negative control (black) cells were cultured in medium with 2 µg/ml doxycycline for 3 days, then 0.05 µM (left) or 0.1 µM of CX-5461 was added to the medium for 0-4 days. Data represent mean ± SD of 3 independent experiments. \*\**P* < 0.01, \*\*\**P* < 0.001. H. Western blots for *TOP2A* (left) and *TOP2B* (right) following CRISPRa induced overexpression. CHP-134 cells were transduced with *TOP2A* or *TOP2B* gRNAs (2 gRNAs for each gene) or no gRNA control (dCas9-VPR). Cell lysates were collected for western blotting. β-Actin was used as a loading control. I. *TOP2A* (red) and *TOP2B* (blue) overexpressing CHP-134 cells and negative control (black) cells were treated with the indicated concentration of CX-5461, etoposide, doxorubicin for 72 hours. Cell viability was determined by MTS assay. Data represent mean ± SD of 3 independent experiments. J. Schematic of *in vitro* topoisomerase activity assay (decatenation assay) workflow. K. Bar plot showing the drug concentration required to inhibit 25% of total DNA decatenation (IC_25_) in an *in vitro* topoisomerase activity assay for a commercial CX-5461 stock, an additional CX-5461 stock that we synthesized, doxorubicin, etoposide and ciprofloxacin. The assay was repeated five times with the error bars representing 95% confidence intervals. Note: we calculated an IC_25_ because CX-5461 did not achieve a reliably measurable IC_50_ within the active drug concentration range for the TOP2A assay (see Fig. S4C). L. Plot showing the relative difference in the concentration of each topoisomerase inhibitor (x-axis) required to inhibit the decatenation of kDNA in the presence of TOP2B relative to TOP2A (y-axis), calculated from the assays performed in panel K. Error bars represent 95% confidence intervals. M. Bar plot showing the drug concentration required to inhibit 50% of total DNA relaxation (IC_50_) for CX-5461, doxorubicin, etoposide and ciprofloxacin. The assay was run twice with repeat runs shown in the left and right panels.

To test this hypothesis, we first performed an shRNA knockdown of *TOP2A* and *TOP2B* using five independent shRNAs in CHP-134 cells. Remarkably, although knockdown of *TOP2A* conferred resistance to etoposide, doxorubicin and CX-5461, knockdown of *TOP2B* only caused resistance to CX-5461 (Fig. 4F-G, *P* < 0.001). The cisplatin-treated negative control was not affected by knockdown of either gene (Fig. 4F). We next used CRISPRa to overexpress both *TOP2A* and *TOP2B* in CHP-134 cells and although overexpression of either gene increased sensitivity to CX-5461, doxorubicin, and etoposide, the effect of *TOP2B* overexpression was greater for CX-5461 (Fig. 4H-I), corroborating the idea that CX-5461 has a strong interaction with TOP2B.

To confirm direct inhibition of topoisomerase II paralogs by CX-5461, we next performed an *in vitro* topoisomerase activity assay, which tests a compound’s ability to inhibit the decatenation of purified catenated kinetoplast DNA (kDNA) plasmids in the presence of purified topoisomerase protein (Fig. 4J). CX-5461 inhibited the activity of both TOP2A and TOP2B— strikingly at concentrations substantially lower than etoposide (Fig. 4K). The negative control ciprofloxacin, which targets type II topoisomerase in prokaryotes and is a fluoroquinolone with structural similarity to CX-5461, had no activity in this assay. By comparing the concentration of CX-5461 required to inhibit TOP2A or TOP2B in this highly controlled *in vitro* system, we directly quantified CX-5461’s preference for TOP2B (Fig. S4C-E). Remarkably, CX-5461 was between and 4-fold more effective at inhibiting TOP2B, compared to TOP2A (*P* = 2.8 × 10^-4^, Fig. 4L). Etoposide and doxorubicin had similar activity against both paralogs. We observed similar trends when we also performed a complementary topoisomerase activity assay to determine CX-5461’s ability to inhibit topoisomerase II-induced relaxation of purified supercoiled DNA, where CX-5461 performed similarly to doxorubicin and etoposide, with a clear preference for TOP2B (Fig. 4M; Fig. S4E). While these multiple orthogonal lines of evidence demonstrate that CX-5461 exhibits classical TOP2 poison activity, our findings are indicative that the primary CX-5461 target is TOP2B.

### Neuroblastoma patient tumors overexpress *TOP2B*

Given our discovery of the mechanism responsible for CX-5461’s robust selective killing of neuroblastoma cell lines, we next sought to determine the potential of CX-5461 to treat neuroblastoma patients. We began by using existing patient genomics datasets to interrogate the *in vivo* activity of the drug target *TOP2B* and to investigate the fidelity of the cell line models used to identify CX-5461. First, it was recently shown that embedding cell lines and patient tumors in a low dimensional latent representation of gene expression data represents a viable strategy to determine whether cancer cell lines recapitulate their tumor-of-origin. We applied this approach^39^ to the DepMap^40^ and GDSC cancer cell lines, which showed that neuroblastoma cell lines were among the most representative of their tumor-of-origin (81% correctly aligned; *P* < 2.2 × 10^-16^, Fig. 5A-B; Fig. S5A), supporting the relevance of the cell line screening data as a model of neuroblastoma patient tumor biology.

**Figure 5.**
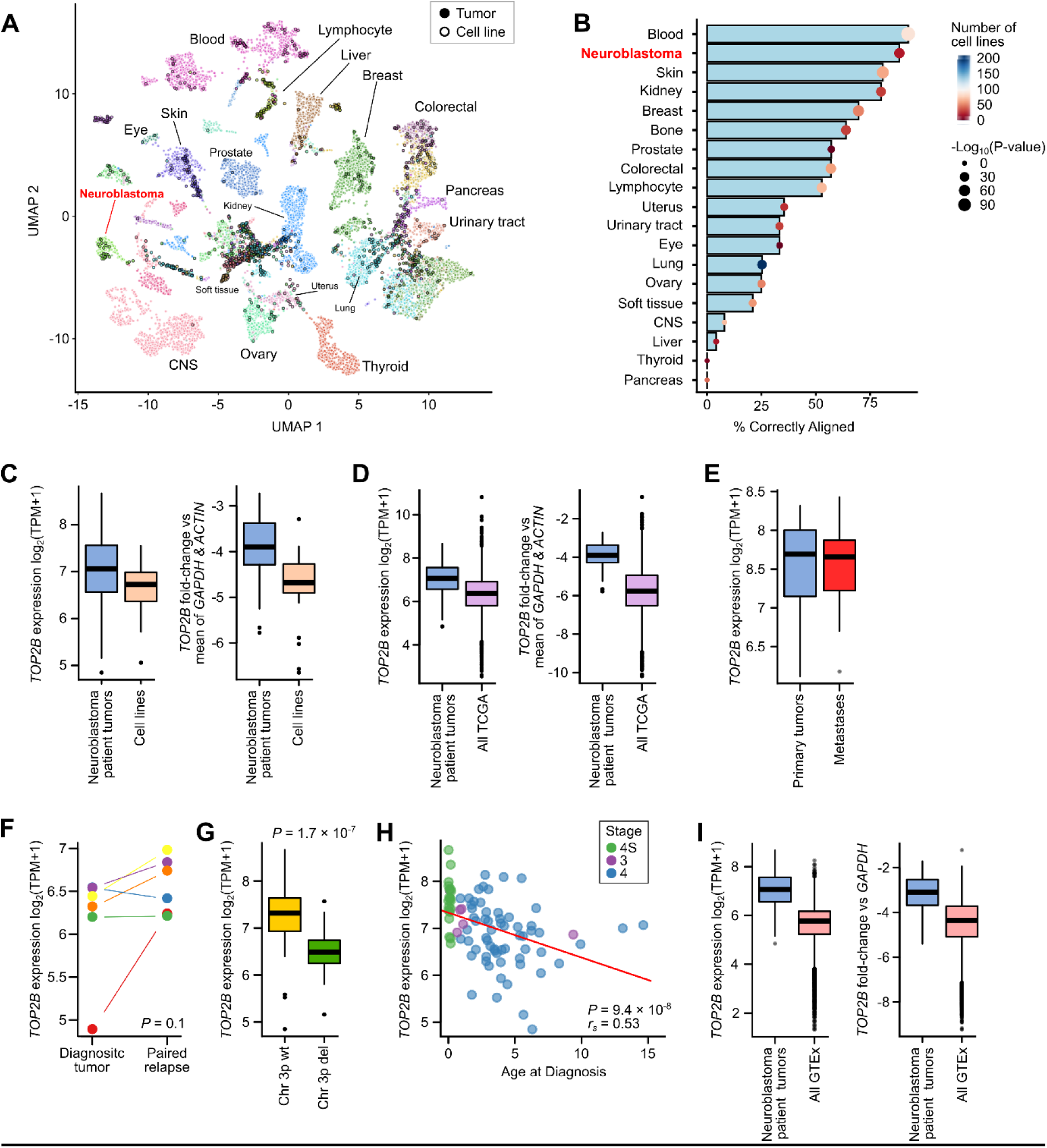
*TOP2B* expression is similar in neuroblastoma patient tumors and cell lines, but it is also expressed in normal cells. A. UMAP representation showing the genome-wide gene-expression-based alignment of all DepMap cell lines with patient tumor gene expression data from TCGA, Treehouse and TARGET, generated using the Celligner tool. The points representing individual cell lines have a black border. Points have been colore by their lineage and clusters have been labeled by tumor lineage. B. The proportion of cell lines from a given lineage that correctly align with the appropriate patient tumor cluster (x-axis). The points have been colored by the number of cell lines in the datasets. The size of the points has been scaled by the *P*-value obtained from a Fisher’s exact test for cluster membership. C. Boxplot of *TOP2B* expression in neuroblastoma cell lines obtained from DepMap RNA-seq vs 88 neuroblastoma patient tumors for log_2_(TPM+1) normalized RNA-seq expression (left) and *TOP2B* expression in these datasets normalized to *ACTIN* and *GAPDH* housekeeping genes (right panel). D. *TOP2B* RNA-seq expression data in 88 neuroblastoma patient tumors compared to all TCGA tumors for log_2_(TPM+1) normalized expression values (left) and *TOP2B* normalized to *ACTIN* and *GAPDH* housekeeping genes (right). E. *TOP2B* log_2_(TPM+1) normalized expression at primary sites and metastatic sites in published RNA-seq data obtained from Rifatbegovic *et al*. F. *TOP2B* log_2_(TPM+1) normalized RNA-seq expression obtained from 6 patients in the TARGET cohort for whom both diagnostic and matched relapse tumor gene expression data were available. G. Boxplot showing *TOP2B* log_2_(TPM+1) normalized RNA-seq expression from 88 neuroblastoma patient tumors with a chromosome 3p deletion compared to the wild-type (wt) copy of chromosome 3p. H. Scatterplot of *TOP2B* log_2_(TPM+1) normalized RNA-seq expression (y-axis) plotted against age (x-axis) for 88 neuroblastoma patient tumors. Patients are colored by disease stage, with stage 4S, 3 and 4 patients colored green, purple and blue respectively. I. *TOP2B* RNA-seq expression data in 88 neuroblastoma patient tumors compared to all GTEx normal tissues for log_2_(TPM+1) normalized expression values (left) and *TOP2B* normalized to *ACTIN* and *GAPDH* housekeeping genes as negative controls (right).

We next explored the *in vivo* activity of *TOP2B* in primary tumors using neuroblastoma patient genomics data. We used an aggregated dataset of 88 diagnostic neuroblastoma tumors where RNA-seq and whole-genome sequencing were available^41^. First, the expression of *TOP2B* in neuroblastoma cell lines was well maintained in patient primary tumors (median expression 104.74 TPM in DepMap neuroblastoma cell lines and 132.4 TPM in our patient tumor cohort; Fig. 5C left). This result was consistent when we normalized against the mean expression of housekeeping genes actin *ACTB (ACTIN)* and *GAPDH* to account for dataset-specific effects (Fig. 5C right). We next assessed the expression of *TOP2B* across TCGA tumor samples. As was true for cell lines, *TOP2B* was more highly expressed in neuroblastoma tumors than other cancers (Fig. 5D, Fig. S5B-C). We also interrogated expression data from relapsed^2^ and metastatic^42^ neuroblastoma tumors, which had paired primary tumor data and we identified no evidence of depletion of *TOP2B* expression in metastases or relapse (Fig. 5E-5F). Together, these data are consistent with the high *TOP2B* expression in neuroblastoma cell lines being maintained in patient tumors.

Notably, the expression of *TOP2B* varies over a 14.57-fold range in these patient tumors (min = 27.8 TPM, max = 405.2 TPM), implying that clinical studies of CX-5461 would benefit from stratifying patients based on the expression of *TOP2B*, the primary drug target. To explore the drivers of these differences, we interrogated the effect of 38 clinical variables and genomic features on *TOP2B* expression. These included somatic mutations, mutation signatures, copy number aberrations, and structural variants. Because of correlations among these features, we used an ElasticNet regression model (α = 0.5) to determine which features were independently predictive of *TOP2B* expression. The model identified 3 predictors; deletion of the 3p arm of chromosome 3 (del(3p)), the q arm of chromosome 11 (del(11q)), and patient age-at-diagnosis (Table S2). *TOP2B* is on chromosome 3p and as expected, del(3p) was associated with a 2-fold decrease in its expression (Fig. 5G). Surprisingly, age-at-diagnosis was also a strong predictor of *TOP2B* expression, with younger patients expressing much higher levels of *TOP2B* (Fig. 5H). *TOP2B* expression was also associated with *MYCN* expression when we conditioned on del(p3) and age-at-diagnosis using multivariate linear regression (*P* = 3.3 × 10^-3^). ChIP-seq data also showed that MYCN is bound to the *TOP2B* promoter (Fig. S5D), consistent with the causal relationship between CX-5461 IC_50_ and *MYCN* expression that we observed in cell lines (Fig. 1G). Overall, these genomics analyses of patient tumors suggest that high TOP2B activity is maintained and may be therapeutically relevant in a subset of neuroblastoma patients. To explore the potential for a CX-5461 therapeutic window in cancer patients, we also assessed the expression of *TOP2B* in normal human tissues using data from the Genotype-Tissue Expression (GTEx) consortium. Although the expression of *TOP2B* was again typically higher in neuroblastoma tumors than normal human tissues (Fig. 5I), there was clear expression across normal tissues, including cardiac tissues and most strikingly some regions of the brain (Fig. S5E-S5F). These genomics analyses provide further confidence that CX-5461’s selective activity against neuroblastoma cell lines could translate to anti-neuroblastoma tumor activity *in vivo. TOP2B*’s expression in normal tissues does, however, highlight potential patient safety concerns given the proposed CX-5461 mechanism-of-action.

### CX-5461 synergizes with inhibitors of ATM, ATR, and TOP1

In the last decade, single-agent clinical trials in neuroblastoma have almost universally failed, even when the rationale for a new compound has been substantive^4^. For such an aggressive disease, identifying synergistic drug combinations is critical. To address this problem, we leveraged our CRISPR screening data, reasoning that druggable proteins whose genetic knockout synergizes with CX-5461 may allow us to narrow the enormous combinatorial search space. For these analyses, we leveraged a previously assembled list of druggable genes^13^. Among the top druggable genes, whose knockout synergizes with CX-5461 in CHP-134 cells were *ATM*, *ATR,* and *TOP1* (Fig. 6A-6B), all genes converging on DNA damage response. Based on our mechanistic work showing the DNA damaging effects of CX-5461 and previous observations of their efficacy in other cancers^43, 44^ we chose to investigate drugs targeting these genes as potential combinations with CX-5461. Thus, we screened CHP-134, IMR-5, BE(2)-M17, and KELLY neuroblastoma cell lines with the combination of CX-5461 and AZD-1390 (ATM inhibitor), AZD-6738 (ATR inhibitor), and SN-38 (the active metabolite of irinotecan; TOP1 inhibitor). We screened each of 12 cell line–drug combination pairs with a comprehensive 12 × 12 matrix of drug concentrations, achieving this scale using high-throughput robotic handling. We estimated regions of synergy using the Zero Interaction Potency (ZIP) model^45^. Encouragingly, we identified regions of synergy at pharmacologically relevant drug concentrations in all 12 cell line-drug combinations (at a fold change of > 1.5 relative to the expectation given an additive model, and *P* < 0.01, Fig. 6C-G; Fig. S6A-L). To confirm the validity of the high-throughput screen, we tested several of these combinations using a conventional MTS assay. In every case, the synergy detected by the MTS assay was consistent with the high-throughput screen, with six representative examples shown in Fig. 6H. Thus, *ATM*, *ATR,* and *TOP1* inhibitors represent promising combinations with CX-5461-targeted TOP2B inhibition to explore in preclinical *in vivo* studies.

**Figure 6.**
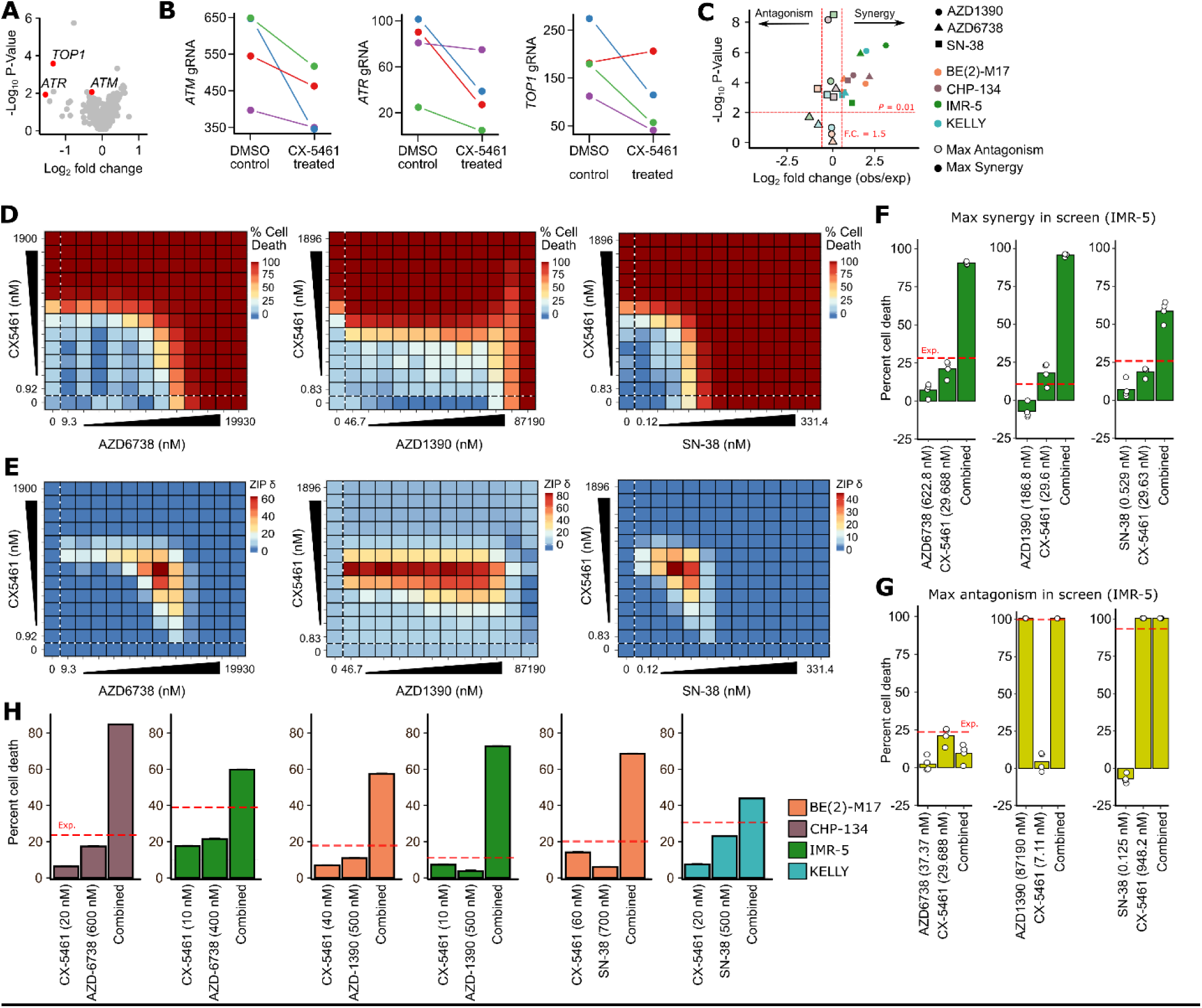
CX-5461 synergizes with inhibitors of ATM, ATR and TOP1. A. Sensitization of CHP-134 cells to CX-5461 following CRISPR knockout of known drug target genes (usin a modified version of the list previously assembled by *Behan et al.*, 2019). B. Normalized guide-RNA (gRNA) read counts (y-axis) for each of four independent guides used to knockout ATM (left), ATR (middle) and TOP1 (right). C. Volcano plot depicting fold-changes (x-axis) and *P*-values (y-axis) at maximum synergy (unbordere shapes) and maximum antagonism (bordered shapes) scores observed in all 12 drug combination screening assays. The shape indicates which drug was combined with CX-5461 and the color of each point indicates the cell line screened. Fold-change is calculated relative to the expectation given additive drug activity and *P*-values were calculated using a one-sample t-test. Thresholds correspond to absolute log_2_(fold change) 1.5 and *P*-value ≤ .01. D. Heatmap matrices of percent cell death in IMR-5 cells conferred by drug-only treatments (values within dotted lines) and combination treatments (all other values) for CX-5461 with AZD6738 (left), AZD139 (middle), and SN-38 (right). Twelve-point doses were used at indicated concentrations with 1:2-fold dilutions for all compounds. Each matrix represents the average of four independent experiments. Data were normalized as a function of percent cell death according to inter-plate controls. E. Heatmap matrices of synergy scores derived from respective cell death values in (D). All synergy was calculated based on the zero interaction potency (ZIP) model, and corresponding scores are shown as δ ZIP according to the original method. Combinations conferring synergy or antagonism correspond to values greater than or less than zero, respectively. F-G. Bar plots of drug combinations that confer the maximum synergy (F) or maximum antagonism (G) scores from their respective synergy calculations. White dots represent four independent experiments corresponding to score maxima. Red dotted lines represent the expected result based on additivity alone. H. Synergy validation with MTS assays for CX-5461 in combination with AZD6738, AZD1390, or SN-38. Colors correspond to cell lines. Data are representative of 3 independent experiments.

### Establishing a pharmacologically relevant *in vivo* murine equivalent dosage for CX-5461

Given the selective activity of CX-5461 against neuroblastoma cell lines, the rational biological mechanism, and the promising synergistic activity with other compounds *in vitro*, we sought to perform an *in vivo* study to test these drug combinations in mice using orthotopic patient-derived xenografts (PDX). To ensure the clinical relevance of our PDX data, we first performed comprehensive *in vivo* preclinical pharmacokinetics studies, comparing CX-5461 plasma area under the concentration-time curve (AUC) and average plasma concentration (Cavg) values to those reported in the available human phase I clinical trial^15^. This is the first time such a study has been reported for this compound. Notably, our PK results demonstrated highly variable CX-5461 plasma concentrations in mice and revealed a much shorter terminal plasma half-life compared to humans (83.3 hours in humans at 250 mg/m2 vs 3.7 hours after 5 mg/kg IV in mice; Fig. S7A-C). This severely undermines the translational relevance of existing preclinical *in vivo* CX-5461 studies in mice, which have not accounted for these differences. To mitigate these issues we devised a low, multiple-dose CX-5461 regimen aimed at maintaining the Cavg above 100 µg/L for 5 days, establishing a dosage of 5 mg/kg once daily by IP injection for 5 days, starting on day 1, with a second 5-day course starting on day 8, every 21 days.

We confirmed on-target drug activity *in vivo* at these pharmacologically relevant dosages using TOP2cc formation and γ-H2AX as pharmacodynamic markers, interrogated using a RADAR assay (Fig. 7A, Fig. S7D), and immunoblotting respectively (Fig. 7B), using PDX tumor tissue harvested from CX-5461 treated mice. Consistent with our *in vitro* observations, we observed no inhibition of RNA-POL I, but saw clear TOP2B trapping and DNA damage in these PDX tumors following CX-5461 treatment (Fig. 7A-C), confirming the *in vivo* relevance of our mechanistic work.

**Figure 7.**
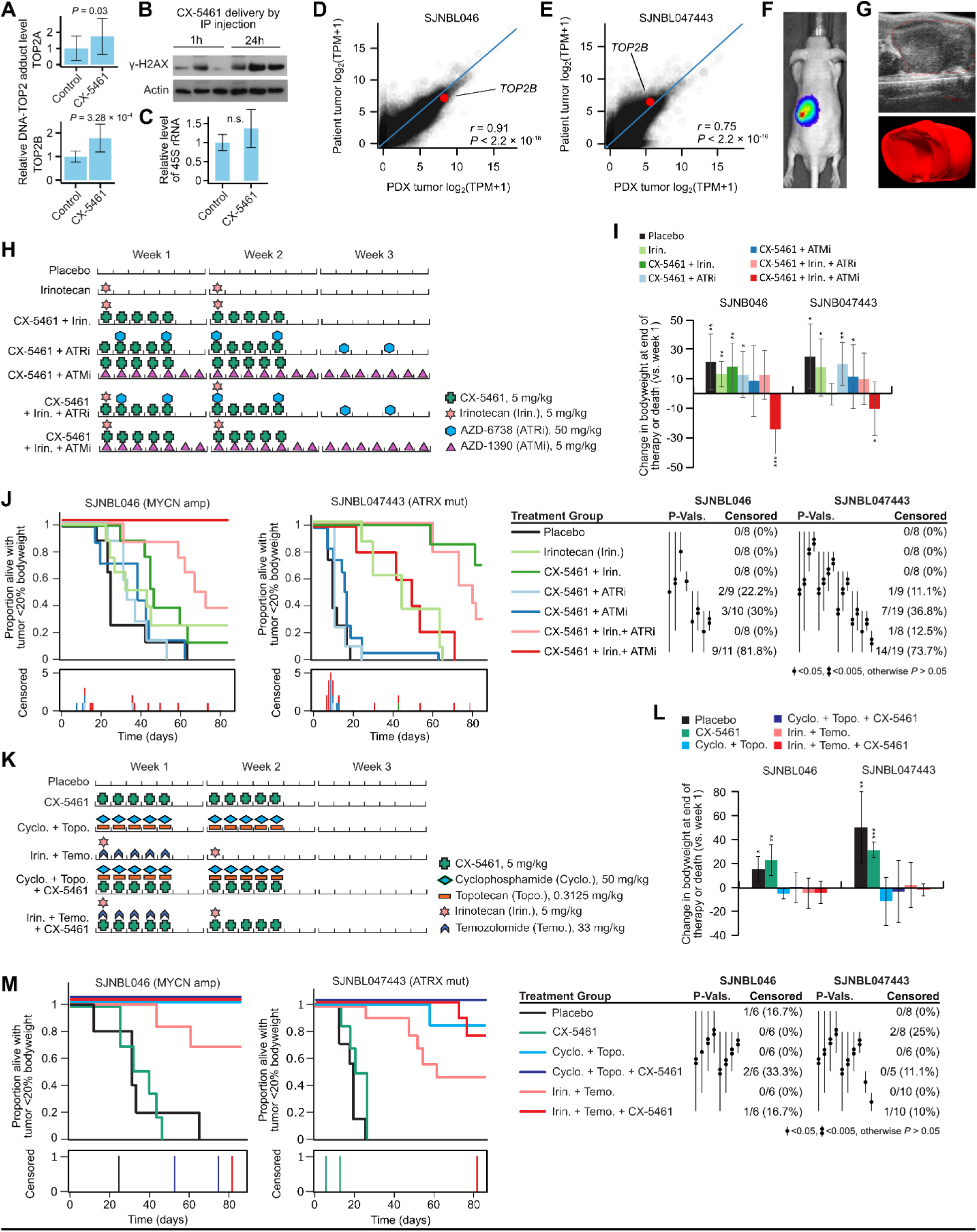
CX-5461 shows promising anti-tumor activity in combination with TOP1 inhibitors *in vivo* using orthotopic PDX mouse models. A. Bar plots showing quantified images from RADAR assay for TOP2-DNA adduct formation (y-axis) for TOP2A (top panel) and TOP2B (bottom panel) in SJNBL046 PDXs harvested from CX-5461 treated mice or control. Tissue for this experiment was obtained from our placebo and CX-5461 group in drug treatment experiments (Fig. 7M). Data represent the mean ± SD of 12 replicates. B. Western blots for γ-H2AX in tumor tissue obtained during PK study. Mice were treated with 25 mg/kg CX-5461 by IP injection and tumors harvested at 1 h or 24 h. β-Actin was used as a loading control. C. Bar plots showing the level of 45s rRNA, measured by qPCR, in the same tissue as Fig. 7A. Data represent the mean ± SD of 12 replicates. n.s., no significant difference. D. Genome-wide gene expression as measured by RNA-seq in *MYCN* amplified patient tumor SJNBL046 (y-axis) and in the PDX derived from this tumor (x-axis). *TOP2B* is highlighted in red. E. Same as (D) but showing data from the ATRX mutant sample SJNBL047443. F-G. Representative image from tumor monitoring using luciferase bioluminescence imaging (F) and ultrasound (G). H. Drug combination schedule depicted for a single 3 weeks cycle in study 1. Mice were treated for 4 cycles. I. Bar plot showing change of bodyweight for each group of mice at the end of therapy or at death (the mice that died before completion of therapy) in study 1. Data represent the mean ± SD. \**P* < 0.05, \*\**P* < 0.01, \*\*\**P* < 0.001. J. Kaplan-Meier curves showing the proportion of mice alive in each treatment group with a tumor burden estimated at <20% of bodyweight, shown for SJNBL046 (left figure) and SJNBL047443 (right figure). The number of mice censored in each treatment group at each time point is shown below. *P*-values were calculated between pairs of curves using a log-rank test and are depicted by a vertical line, with the number of dots corresponding to the level of significance; comparisons yielding *P* > 0.05 are not shown. K. Drug combination schedule depicted for a single 3 weeks cycle in study 2. Mice were treated for 4 cycles. L. Like (I) but showing bodyweights for study 2. M. Like (J) but showing the results of study 2.

### CX-5461 improves survival *in vivo* when combined with TOP1 inhibitors in orthotopic PDX mouse models

We performed our *in vivo* drug treatments using two luciferase-labeled PDXs—SJNBL046, a *MYCN* amplified model from a 2-year-old male, and SJNBL047443, an ATRX mutant model from a 12-year-old male. Both PDXs were from pre-treated neuroblastoma tumors, with SJNBL046 surgically resected after induction chemotherapy and SJNBL047443 resected at disease recurrence. We used RNA-seq to compare genome-wide gene expression in these PDXs to their patient tumor of origin and the expression profiles were well maintained (*r* = 0.91, *P* < 2.2 × 10^-16^ for SJNBL046; *r* = 0.75, *P* < 2.2 × 10^-16^ for SJNBL047443; Fig. 7D-E); *TOP2B* expression was also maintained, suggesting these PDXs are a relevant disease model. We implanted the cells by ultrasound-guided injection into the orthotopic para-adrenal region of athymic-nude mice, and we monitored tumors using bioluminescent imaging and weekly ultrasound (Fig. 7F-G; Fig. S7E-F).

We first tested 2-drug combinations of CX-5461 with pharmacologically relevant dosages of AZD-1390 (ATMi), AZD-6738 (ATRi), and the liposomal formulation of the TOP1 inhibitor irinotecan. We also tested 3-drug combinations involving CX-5461, irinotecan, and the ATMi or ATRi (Fig. 7H). Combinations involving CX-5461 + ATMi had clear toxicity and were associated with lower body weight (Fig. 7I; Fig. S7G). Indeed, for the CX-5461 + ATMi two-drug combination, 30% (MYCN amplified group) and 36.8% (ATRX mutant group) of mice were withdrawn (censored) from the study, primarily due to toxicity; this increased to 81.8% and 73.7% with the addition of irinotecan (Fig. 7J, Tables S3-S4). The combinations involving ATRi lacked efficacy *in vivo*, with the two-drug combination of CX-5461 + ATRi showing no activity beyond placebo, and no increase in efficacy was observed with the addition of ATRi to CX-5461 + irinotecan (Fig. 7J). However, the dual treatment of CX-5461 with the TOP1 inhibitor irinotecan was effective, resulting in improved outcomes relative to placebo for both PDXs (*P* = 0.03 and *P* = 9.4 × 10^-5^ for MYCN and ATRX PDXs respectively; Tables S5-S6). Importantly, no mice were censored due to toxicity.

We designed a second study to further explore the combination of CX-5461 with TOP1 inhibitors, combining CX-5461 with two 2-drug salvage combinations used in relapsed and refractory neuroblastoma, which have been used as backbones for incorporating new agents in clinical trials: irinotecan + temozolomide and topotecan + cyclophosphamide (Fig. 7K; Table S7-S8)^46, 47^. The therapies were reasonably tolerated with minimal loss of body weight (Fig. 7L; Fig. S7H) or censoring (Figure 7M). For both regimens, the addition of CX-5461 improved outcome in every group in both PDXs, clearest in the irinotecan + temozolomide combination in the SJNBL047443 PDXs (*P* = 0.05; Tables S9-S10). Thus, in all six independent experiments, CX-5461 improved survival when added to a TOP1 inhibitor (*P* = 0.01). The CX-5461 treated mice also survived longer following cessation of therapy in all groups (mean 71 to 83.75 days and 90.8 to 144 days for the addition of CX-5461 to cyclophosphamide and topotecan, 21.75 to 24.8 days and 4 to 18.8 days for the addition of CX-5461 to irinotecan and temozolomide for the MYCN and ATRX mutant PDX groups respectively; Table S7-S8).

Overall, our mouse studies suggest that combinations of CX-5461 with TOP1 inhibitors are synergistic *in vivo* and have clinical potential in neuroblastoma. However, despite the anti-tumor activity, our mechanistic work raises the possibility of late-emerging adverse events when TOP2B is targeted, which needs to be further understood before this combination is applied in a pediatric patient cohort.

## DISCUSSION

We used large-scale preclinical drug screening data to identify CX-5461 as exhibiting a selective and reproducible cytotoxic effect in high-risk neuroblastoma and we discovered that the primary target of CX-5461 at pharmacologically relevant concentrations is TOP2B. Because of the functionally unique role of TOP2B, this finding completely alters the clinical trajectory of CX-5461 in terms of patient stratification, rational combinations, potential resistance mechanisms, and safety profile.

Evidence that CX-5461 inhibits the transcription of rRNA has fueled previous conclusions that CX-5461 directly targets the RNA-POL I machinery^48^. Several studies have implicated CX-5461 as a binder of DNA G-quadruplexes *in vitro*^16, 49^, and a single systems biology study argued that CX-5461 targets TOP2A^18^ (TOP2B was not mentioned in ref. 18). However, none of these previously proposed mechanisms of CX-5461 unify the wide range of observations made on this compound—nor identified its molecular target. The activity of CX-5461 as a TOP2B inhibitor is consistent with numerous previous observations, including CX-5461’s DNA damaging effects^16^ and its tendency to inhibit RNA-POL I-mediated rRNA transcription^25^, even in non-cycling cells that do not express *TOP2A*^37^. Multiple previous studies have shown that topoisomerases play a key role in resolving DNA topological issues to maintain active transcription^50^, including at the rRNA loci^51^ and inhibition of topoisomerases (and treatment with topoisomerase inhibitors) has been well established to cause inhibition of RNA-POL I^52, 53^. Additionally, TOP2B has been shown to localize to the rRNA locus, where RNA-POL I activity occurs^54^. Thus, our study leads us to a parsimonious model, able to explain the capabilities of CX-5461 to induce DNA damage and inhibit rDNA transcription: specifically, that the molecular target of CX-5461 is TOP2B, and that previously observed inhibitory effects of CX-5461 on RNA-POL I are a consequence of topoisomerase inhibition.

Interestingly *TOP2B* knockout mice die shortly after birth due to defects in the peripheral nervous system occurring during the late stages of development^55^, suggesting *TOP2B*’s relevance as a drug target in neuroblastoma partially stems from the tumor’s developmental cell-of-origin. We showed that *TOP2B* remains overexpressed in neuroblastoma, particularly younger and neonatal patients and that *TOP2B* expression is driven higher by *MYCN* overexpression, explaining the selective activity of CX-5461 against neuroblastoma cells. However, even though we showed anti-tumor activity in combination with TOP1 inhibitors *in vivo* in orthotopic PDX models at pharmacologically relevant dosages, the clinical translation of CX-5461 in neuroblastoma faces an unexpected obstacle—TOP2B as CX-5461’s primary target raises major previously unanticipated safety concerns. Existing TOP2A inhibitors can cause serious *late-emerging* side effects, which would not be observable in a typical preclinical study, nor the short-duration phase I clinical trial. For example, etoposide leads to therapy-induced AML 2-3 years after treatment in about 1% of patients, and doxorubicin causes (frequently fatal) cardiotoxicity in about 11% of patients, both of which are caused by off-target interactions with TOP2B^20, 21^. Consistent with this, our analysis of GTEx data showed that *TOP2B* is broadly and highly expressed in normal human tissues, which is not true of *TOP2A*. CX-5461 has recently been shown to be mutagenic in a *C. elegans* model^56^, contradicting earlier claims that it is not genotoxic^25^. Thus, our finding that CX-5461 primarily targets TOP2B provokes serious safety concerns, and patients enrolled in ongoing CX-5461 phase II trials should be very closely monitored for these late-emerging *TOP2B*-related adverse events. *TOP2B* is also particularly highly expressed in the brain, thus blood-brain barrier penetration should be formally assessed.

Our analysis of TCGA data showed that subsets of both breast and hematological cancer patients (the diseases of ongoing trials) also overexpress *TOP2B*, most strikingly in some leukemias. Our identification of CX-5461’s primary target will facilitate patient stratification in these trials, which along with rational drug combinations will markedly improve the chances of successful phase II trials in adult cancers, if safety concerns can be addressed. Indeed, given the concerns surrounding the safety of TOP2B as a primary drug target, achieving promising results in an adult cohort will likely be a prerequisite for a clinical study in children, where the barriers to new clinical trials are much higher due to the limited and highly vulnerable patient population.

## ACKNOWLEDGEMENTS

PG is supported by the NIH, including an R00 award from NHGRI (5R00HG009679-03) and an R35 award from NIGMS (1R35GM138293-01). The content is solely the responsibility of the authors and does not necessarily represent the official views of the National Institutes of Health. PG also receives funding from ALSAC. The authors acknowledge Dr. Jinghui Zhang and Dr. Alex Gout for critical review of the manuscript, and Melissa D. Johnson, Thomas Confer, and Walter J. Akers for performing the *in vivo* imaging work.

## AUTHOR CONTRIBUTIONS

P.G. conceived and directed the study. M.P. performed the experimental work. P.G. and M.P. wrote the manuscript. P.G. and J.E. supervised the experimental work. C.W., R.C., M.S., A.Z., S.B., D.M., M.P., and P.G. performed computational work. J.B. and S.B. assisted with topoisomerase activity assays. J.L. and S.C. performed drug screening, with C.W. analyzing the data. Y.W. performed pharmacokinetic and bioanalytical assays, supervised by B.B.F. K.B.B., B.G., P.A. and E.S. performed the PDX experiments, directed by E.S. P.C. and S.M.P.M assisted with CRISPR screening. J.B., S.N. and M.J.P. performed additional supportive experimental work. R.K., M.M.B, M.A.D, J.S., T.C., L.A.G & S.B provided resources, insights and supervised additional parts of the study. All authors edited the final manuscript.

## METHODS

### Cell Lines and Cell Culture Conditions

CHP-134, KELLY, and SK-N-SH cells were purchased from Sigma (MilliporeSigma, USA). BE(2)-M17, SK-N_FI, RS4-11, H2452, and 293T cells were purchased from ATCC. IMR-5 cell line was a gift from the Cellular Screening Center at the University of Chicago. IMR-5 was cultured in Eagle’s Minimum Essential Medium (EMEM, ATCC catalog# 30-2003) supplemented with fetal bovine serum (FBS, ATCC® 30-2020™) to a final concentration of 10%. The rest of the cell lines were cultured under conditions recommended by the vendors. All cells are cultured at 37°C with 5% CO_2._ No cell lines used in this study were found in the database of commonly misidentified cell lines that is maintained by ICLAC and NCBI Biosample. Cell lines were tested negative for mycoplasma using the MycoAlert mycoplasma detection kit (Lonza, LT07-118).

### Mice and Housing Conditions

Athymic nude immunodeficient mice were purchased from Charles River (strain code 553). This study was carried out in strict accordance with the recommendations in the Guide to Care and Use of Laboratory Animals of the National Institute of Health. The protocol was approved by the Institutional Animal Care and Use Committee at St. Jude Children’s Research Hospital. All efforts were made to minimize suffering. All mice were housed per approved IACUC protocols. Animals were housed on a 12-12 light cycle (light on 6 am off 6 pm) and provided food and water *ad libitum*.

### Transfection and transduction

Inducible TRIPZ lentiviral shRNA of human TOP2A, TOP2B, and negative control shRNA plasmids were purchased from Dharmacon (Horizon Discovery, USA. TOP2A shRNA clone numbers are RHS4696-200686220, RHS4696-200769079. TOP2B clone numbers are RHS4696-200690242, RHS4696-200702560, RHS4696-200755958). MYCN shRNAs were ordered from Transomic Technologies (Cat# TLHSU2300-4613. Clone numbers are ULTRA-3327035, ULTRA-3327037, ULTRA-3475807). Lentiviral envelope and packing plasmids pMD2.G and psPAX2 were ordered from Addgene (pMD2.G #12259, psPAX2 #12260). To generate lentiviral particles, 293T cells were seeded in a 6-well plate one day before transfection. For one well of a 6-well plate, cells were co-transfected with 0.56 µg pMD2.G, 0.83 µg psPAX2, and 1.1 µg shRNA plasmid, using Lipofectamine 3000 (ThermoFisher, USA. Catalog# L3000008). Cell culture medium with lentivirus were collected three times during the time period 36 h to 72 h after transfection. CHP-134 cells were infected two to three times during 24 h. Two days after infection, 0.5 µg/ml puromycin was added to the culture medium to select infected positive cells. 2 µg/ml doxycycline was used to induce shRNA expression.

CRISPR activation sgRNA plasmids for TOP2A (Cat# GSGH11887-247255376, GSGH11887-247255378) and TOP2B (Cat# GSGH11887-247255386, GSGH11887-247255388), and plasmids for dCas9 expression (Cat# CAS11914) were ordered from Horizon Discovery. Lentiviral particles were generated with co-transfection with pMD2.G and psPAX2 in 293T cells. CHP-134 cells were infected with dCas9 lentivirus and dCas9-expressing CHP-134 cells were selected with 7 µg/ml blasticidin. The virus with TOP2A or TOP2B CRISPR activation sgRNAs were used to infect the blasticidin-resistant cells. Positive cells with TOP2A or TOP2B sgRNAs were selected with 0.5 µg/ml puromycin.

### RNA extraction and quantitative real-time RT-PCR

Total RNA was extracted using Trizol (Invitrogen) according to the manufacturer’s protocol. cDNA was synthesized using qScript cDNA Synthesis Kit (Quanta Biosciences, catalog# 95047). SYBR Green PCR Master Mix (Quanta Biosciences, catalog# 95072-012) was used for the quantitative real-time RT-PCR, which was performed on the CFX96 real-time PCR detection system (Biorad). Experimental results were normalized to β-Actin. The primers for 45S pre rRNA internal transcribed spacer region and 18s rRNA were previously described ^16^ (45S-F 5’-GCCTTCTCTAGCGATCTGAGAG-3’, 45S-R 5’-CCATAACGGAGGCAGAGACA-3’; 18S-F 5’-AAACGGCTACCACATCCA-3’, 18S-R 5’-CCTCCAATGGATCCTCGT-3’). 28S-rRNA primers were previously described ^57^ (28S-F 5-GGGTGGTAAACTCCATCTAAGG-3, 28S-R 5′GCCCTCTTGAACTCTCTCTTC-3′). β-Actin-F 5’-CCAACCGCGAGAAGATGA-3’, β-Actin-R 5’-CCAGAGGCGTACAGGGATAG-3’

### Flow cytometry

For apoptosis, cells were treated in 6-well plates. After incubation for the time needed, cell culture medium and cells were collected and washed in cold phosphate-buffered saline (PBS). The cells were then resuspended in annexin-binding buffer and stained with Alexa Fluor 488 conjugated annexin V (Thermo Fisher, catalog# A13201) and 1 µg/ml Propidium iodide (PI) for flow cytometry analysis. For cell cycle analysis, cells were fixed with cold 70% ethanol in water. After 2× washes in PBS, cells were treated with 100 µg/ml RNase A and stained with 50 μg/ml PI for flow cytometry analysis.

### MTS and CellTiter-Glo assays

Cells were seeded in 96-well plates and treated with compounds as needed. Premixed MTS reagent (Abcam, ab197010) was added to the medium in a 96-well plate to a final concentration of 10%. Then the clear bottom plate was incubated in a cell culture incubator for 0.5 to 4 hours and absorbance was measured at 490nm. For CellTiter-Glo (Promega, G9241), reagent was added to the white bottom plate to a final concentration of 50%. The plate was then shaken for 2 min to induce cell lysis. After incubation at room temperature for 10 minutes, the luminescence was recorded.

### Western blotting and band depletion assay

Generally, cells were washed with cold PBS and lysed in RIPA buffer (1% sodium deoxycholate, 0.1% SDS, 1% Triton X-100, 10 mM Tris [pH 8.0], and 150 mM NaCl) in cell culture plates. RIPA buffer was supplemented with Protease Inhibitor Cocktail (Sigma, #5892970001) and Phosphatase Inhibitor (Thermo Fisher, #A32957). For cleaved PARP detection, cell culture medium was also collected. Protein concentration was determined by BCA protein assay kit (Thermo Fisher, catalog# 23225). After boiled in 1× LDS sample buffer (Thermo Fisher, #NP0007), proteins were separated by SDS-PAGE and transferred to PVDF membranes. After blocking in 5% non-fat milk in TBS with 0.1% Tween-20, membranes were probed with primary antibodies and horseradish-peroxidase-conjugated secondary antibodies. The signal was visualized by Western Lightning Plus-ECL (PerkinElmer). Band-depletion assay was performed as previously described with slight modification ^33^. Treated cells were lysed in culture plates with 1× Laemmli sample buffer (2% SDS, 10% glycerol, 0.002% bromphenol blue, 0.0625M Tris-Cl (pH 6.8), 10% 2-mercaptoethanol). Samples were boiled and 10 µl of each sample was separated by SDS-PAGE. The antibodies used for western blotting in this study are β-Actin (Sigma, #A1978), cleaved PARP (Cells Signaling, #5625), TOP1 (Santa Cruz, #sc-271285), TOP2A (Santa Cruz, #sc-365916), TOP2B (BD Biosciences, #611492), FLAG-Tag (Cell Signaling, #2368), P53 (Santa Cruz, #sc-126), phospho-P53 S15 (Cell signaling, #9284), pCHK1 S345 (Cell Signaling, #2348), CHK1 (Santa Cruz, #sc-8408), pCHK2 T68 (Cell Signaling, #2661), CHK2 (Santa Cruz, #sc-17747), HRP-linked anti-rabbit IgG (Cell Signaling, #7074), HRP-linked anti-mouse IgG (Cell Signaling, #7076).

### Immunofluorescence, EU and EdU incorporation

Cells were seeded on poly-D-lysine coated coverslips before treatment. Treated cells were fixed in 4% formaldehyde for 15-20 minutes and permeabilized with 0.3% Triton X-100 for 10 min at room temperature. For γ-H2AX, 53BP1, and RPA staining, cells were blocked in 5% FBS/PBS for 1 h at room temperature, then incubated with primary antibody diluted in 1% FBS/PBS overnight at 4°C. Fluorochrome-conjugated secondary antibodies were incubated for 1 h at room temperature. For G-quadruplex detection, RNA in fixed cells was digested by incubating cells in 0.5 mg/ml RNase A for 1 h at 37°C. Then block coverslips with 0.5% goat serum in PBST (PBS + 0.1% Tween20) for 1 h at 37°C. Primary antibody 1H6 (Millipore, # MABE1126) was diluted in 0.5% goat serum/PBST and incubate with coverslips at 37°C for 1 hr or at 4°C overnight. Alexa Fluor 488 conjugated secondary antibody was diluted in 0.5% goat serum/PBST and incubate at room temperature for 1 h. ProLong® Gold Antifade Reagent with DAPI (Cell Signaling, #8961) was used as coverslip mountant. Coverslips were mounted for 24 h at room temperature and subjected to fluorescence microscopy.

For ethynyl uridine (EU) and 5-ethynyl-2’-deoxyuridine (EdU) incorporation assay, Click-iT™ RNA Alexa Fluor™ 594 Imaging Kit (Thermo Fisher, #C10330) and Click-iT™ EdU Alexa Fluor™ 488 Imaging Kit (Thermo Fisher, #C10337) were used. Experiments were carried out following the manufacture’s protocols. Briefly, cells were cultured in the medium with EU or EdU for 30 min to 3 h. After fixation and permeabilization, cells were incubated for 30 minutes at room temperature in EU or EdU Click-iT® reaction cocktail and then washed in Click iT® reaction rinse buffer (for EU) or 3% BSA in PBS (for EdU). If needed, cells were processed for additional staining as regular immunofluorescence.

Images were taken with the Nikon C2 confocal microscope and analyzed with NIS-Elements Viewer imaging software. γ-H2AX signal were quantified and analyzed using ImageJ. The antibodies used for immunofluorescence are γ-H2AX (Cell Signaling, #9718; Millipore, # 05-636), 53BP1 (Abcam, ab36823), RPA (Millipore, # MABE285), Alexa fluor 488 conjugated goat anti-mouse IgG (Thermo Fisher, #A-11001), Alexa fluor® 594 conjugate anti-rabbit IgG (Cell Signaling, 8889).

### Comet assay

Comet assay was performed with the Comet Assay Kit (Abcam, ab238544), following the protocol from the manufacture. In brief, cells were collected and counted, then mixed with Comet Agarose at 1/10 ratio (v/v). The cell and agarose mixture was immediately transferred onto an agarose coated glass slide. After solidified at 4 °C for 15 min, the slide was incubated in pre-chilled Lysis Buffer for 30-60 mins at 4°C in the dark, then in pre-chilled Alkaline Solution for 30 mins at 4°C in the dark. The slide was then subjected to electrophoresis in Alkaline Solution. After fixed in 70% ethanol, DNA was stained with Vista Green DNA Dye. Slides were viewed and imaged with Nikon C2 microscope at 10× magnification. Images were analyzed with OpenComet tool.

### RADAR assay

The assay was performed as previously described ^34^. Treated cells were lysed with MB buffer (6M guanidinium isothiocyanate, 10 mM Tris-HCl, pH 6.5, 20 mM EDTA, 4% Triton X100, 1% Sarkosyl, and 1% dithiothreitol). Nucleic acids were precipitated by adding 100% ethanol to the cell lysate to a final concentration of 33% (1/2 volume of the cell lysate). After 5 min of incubation at −20 °C, nucleic acids were collected by centrifugation. Pellet was washed with 75% ethanol and dissolved in freshly made 8 mM NaOH solution. DNA concentration was determined by NanoDrop 1000 spectrophotometer. 1 µg DNA from each sample was transferred to a nitrocellulose membrane using the Bio-Dot SF microfiltration apparatus (Bio-Rad, #170-6542). The membrane was blocked in 5% non-fat milk in TBS with 0.1% Tween-20, then probed with primary antibodies and horseradish-peroxidase-conjugated secondary antibodies. The signal was visualized by Western Lightning Plus-ECL (PerkinElmer). Antibodies used were listed in section ‘Western blotting and band depletion assays’.

### CRISPR screen

CHP-134 cells were transduced with lentiCas9-Blast lentivirus. Media with 7 µg/ml blasticidin was added 2 days later. After 5 days of selection, a single cell was seeded to each well of a 96-well plate by flow cytometry single-cell sorting. Single clones were picked out after 3 weeks and Cas9 expression level in each clone was determined by western blotting with Cas9 antibody (Cell Signaling, #14697). Cas9 activity was tested using the CRISPRtest™ Functional Cas9 Activity Kit for Human Cells (Cellecta, #CRTEST). The clone with normal viability and high Cas9 activity (> 95%) was used for the following screening.

CHP-134 Cas9 cells were transduced at day 0 with lentiviral Brunello pooled library ^58^ at a low MOI to make the transduction efficiency ∼30% ^59^. 0.5 µg/ml puromycin was used to select transductants after 2 days of transduction (day 2). Cells were subcultured on day 4. On day 6, cells were split into 2 groups, with 500 representatives for each gRNA in each group. On the next day (day 7), one group of cells were treated with 0.04 µM CX-5461, DMSO was added to the other group. All cells were collected for DNA extraction after 3 days (day 10).

The Brunello sgRNA pooled plasmid library (Addgene #73178) and lentiCas9-Blast plasmid (Addgene #52962) were expanded following manufacturer’s guidelines at The Center for Advanced Genome Engineering at St. Jude Children’s Research Hospital. Lentiviral particles were generated by the St. Jude Vector Development and Production Core. Genomic DNA was extracted using Nucleospin Blood XL kit (Macherey-Nagel cat# 740950) and eluted into 1ml elution buffer. gDNA samples were amplified for next-generation sequencing at the St. Jude Center for Advanced Genome Engineering using primer sets and reagent recommended by the Broad Genetic Perturbation Platform (https://portals.broadinstitute.org/gpp/public/). Samples were sequenced by the St. Jude Hartwell Center for Bioinformatics and Biotechnology and resulting sequencing data analyzed using MAGeCK ^60^.

### Animal experiments

Isolation of SJNBL046_X Cells: The tumor material was collected from a male patient diagnosed with MYCN-amplified neuroblastoma at St. Jude Children’s Research Hospital, after informed consent and in agreement with local institutional ethical regulations. Tumor tissue was initially processed within 2 hours of surgical resection into a single cell suspension by enzymatic dissociation and injected into the para-adrenal region of NSG mice using ultrasound guidance as described below. Tumor growth was monitored with ultrasound and manual palpation. After tumor development, the tumor was harvested and passaged into athymic nude mice using the same dissociation and ultrasound implantation techniques.

Isolation of SJNBL047443_X Cells: The tumor material was collected from a male patient diagnosed with ATRX-mutated neuroblastoma at St. Jude Children’s Research Hospital, after informed consent and in agreement with local institutional ethical regulations. Tumor tissue was initially processed within 2 hours of surgical resection and implanted into the flank of an NSG mouse. After tumor development, the tumor was harvested, processed into a single cell suspension by enzymatic dissociation, and passaged into athymic nude in the para-adrenal location using ultrasound guidance.

Orthotopic Ultrasound-Guided Para-adrenal Xenograft Implantation: All ultrasound procedures were performed using the VEVO 3100 high-frequency ultrasound equipped with MS-MX250 24 MHz, MX550D 55 MHz, and MX700 70 MHz transducers (Fujifilm-Visualzonics, Toronto, ON, Canada). Cells were suspended in Matrigel (BD Worldwide, Cat#354234) at a concentration of 2×10^4^ cells per microliter and placed on ice. Anesthetized recipient athymic nude mice (isoflurane 2-3% in 100% oxygen and delivered at 2 liters/min) were placed laterally on the heated platform in right lateral recumbency. To provide a channel for delivery of the implant, a 22 gauge catheter (BD Worldwide, Cat# 381423) was gently inserted through the skin and back muscle into the para-adrenal region, the stylet removed, and the hub clipped off. A chilled Hamilton syringe fitted with a 27 gauge needle (1.5 inches) and loaded with 10 µl of the cell suspension was inserted using a micromanipulator through the catheter and the tip positioned between the kidney and adrenal gland via ultrasound guidance. The cell suspension was injected into the region and the needle left in place for 30 seconds to permit the Matrigel component to set before retracting the needle and catheter.

Statistical analysis: P-values for pairwise comparisons of all survival curves within each experiment were calculated using a log-rank test, implemented in *survdiff()* function in the R package “survival”. Censoring plots and survival plots were created in R and Inkscape.

#### Study 1

Neuroblastoma orthotopic xenografts were created by injecting luciferase labeled SJNBL046_X and SJNBL047443_X cells into recipient athymic nude mice using the para-adrenal injection technique previously described. Mice were screened weekly by ultrasound for tumor volume measurements. Mice were enrolled in the study when the tumor volume was the size of the adrenal gland (8mm^3^) or above and chemotherapy was started the following Monday.

SJNBL046_X mice (n = 64) and SJNBL047443_X mice (n=71) were randomized to 9 treatment groups: liposomal irinotecan (Irin.), Irin. + CX-5461, Irin. + CX-5461 + AZD6738, Irin. + CX-5461 + AZD1390, CX-5461 + AZD1390, CX-5461 + AZD6738, and placebo. The following doses were used:

| TREATMENT GROUP | DOSING |
| --- | --- |
| Irin. | 5 mg/kg once daily on days 1 and 8 |
| Irin. + CX-5461 | Irin. 5 mg/kg once daily on days 1 and 8, CX-5461 5 mg/kg once daily on days 1-5 and 8-12 |
| Irin. + CX-5461 + AZD6738 | Irin. 5 mg/kg once daily on days 1 and 8, CX-5461 5 mg/kg once daily on days 1-5 and 8-12, AZD6738 50 mg/kg once daily on days 2, 5, 9, 12, 16, and 19 |
| Irin. + CX-5461 + AZD1390 | Irin. 5 mg/kg once daily on days 1 and 8, CX-5461 5 mg/kg once daily on days 1-5 and 8-12, AZD1390 5 mg/kg once daily on days 1-21 |
| CX-5461 + AZD1390 | CX-5461 5 mg/kg once daily on days 1-5 and 8-12, |
|  | AZD1390 5 mg/kg once daily on days 1-21 |
| CX-5461 + AZD6738 | CX-5461 5 mg/kg once daily on days 1-5 and 8-12,<br>AZD6738 50 mg/kg once daily on days 2, 5, 9, 12, 16, and<br>19 |
| PLACEBO | vehicle control (no chemotherapy) |

#### Study 2

Neuroblastoma orthotopic xenografts were created by injecting luciferase labeled SJNBL046_X and SJNBL047443_X cells into recipient athymic nude mice using the para-adrenal injection technique previously described. Mice were screened weekly by ultrasound and tumor volume was measured. Mice were enrolled in the study when the tumor volume was the size of the adrenal gland (8mm^3^) or above and chemotherapy was started the following Monday. Mice received 4 courses of chemotherapy and ultrasound tumor volume was monitored between courses and at the end of therapy. In addition to ultrasound, bioluminescence imaging was performed weekly. Mice were monitored daily while receiving chemotherapy and removed from the study and euthanized if found to be moribund at any time. Disease response was assigned according to tumor size. Mice that did not have a tumor at the end of the study by ultrasound and necropsy were classified as complete response. Mice that completed all 4 courses of therapy and had a measurable tumor less than 20% of the bodyweight were classified as a partial response. Mice with a tumor burden greater than 20% of the bodyweight at any point were removed from the study and classified as progressive disease.

SJNBL046_X mice (n = 32) and SJNBL047443_X mice (n=44) were randomized to 6 treatment groups: CX-5461, liposomal irinotecan (Irin.) + temozolomide (Temo.), cyclophosphamide (Cyclo.) + topotecan (Topo.), Cyclo. + Topo. + CX-5461, Irin. + Temo. + CX-5461, and placebo. The following doses were used:

| TREATMENT GROUP | DOSING |
| --- | --- |
| CX-5461 | 5 mg/kg once daily on days 1-5 and 8-12 |
| Irin. + Temo. | Irin. 5 mg/kg once daily on days 1 and 8, Temo. 33mg/kg once daily on days 1-5 |
| Cyclo. + Topo. | Cyclo. 50 mg/kg once daily on days 1-5, Topo. 0.3125 mg/kg once daily on days 1-5 |
| Cyclo. + Topo. + CX-5461 | Cyclo. 50 mg/kg once daily on days 1-5, Topo. 0.3125 mg/kg once daily on days 1-5, CX-5461 5 mg/kg once daily on days 1-5 and 8-12 |
| Irin. + Temo. + CX-5461 | Irin. 5 mg/kg once daily on days 1 and 8, Temo. 33mg/kg once daily on days 1-5, CX-5461 5 mg/kg once daily on days 1-5 and 8-12 |
| PLACEBO | vehicle control (no chemotherapy) |

Abdominal Ultrasound and Tumor Volume Acquisition: The Vevo 3100 system was used for ultrasound measurements of orthotopic neuroblastoma xenografts. Mice were anesthetized with isoflurane (2%-3% in 100% oxygen) and positioned on the heated platform. The abdomen was coated with ultrasound gel (Aquasonics, Parker Labs, Fairfield, NJ) and a MX250 or MX550 transducer used for 3D B-mode image acquisition. For volumetrics, 2D image slices were acquired in the coronal plane covering a 20-30 mm range with a step-size of 0.076mm. All post-processing of data was performed using Vevo LAB 3.2 software. Briefly, tumor margins were manually segmented at 5-10 slice intervals, closed at the cranial and caudal ends and reconstruction was performed to identify final volumes.

Bioluminescence Imaging and Quantification: Mice were anesthetized with isoflurane (2-3% in 100% oxygen) and bioluminescence images acquired with an IVIS Spectrum (Perkin Elmer, Waltham, MA) 5 minutes after intraperitoneal injection of Potassium D-Luciferin solution (Perkin Elmer, 3 mg/mouse). Luminescence values for all xenografts were quantified as the average radiance (photons/s/cm2/sr) from identical regions of interest (ROI) encompassing the entire abdomen using Living Image 4.7 (Perkin Elmer).

### Preclinical pharmacokinetic studies

Previously published CX-5461 dosages in mice have ranged from 12-50 mg/kg orally (PO) or by intraperitoneal (IP) injection, from once weekly to continuous 5-day on, 2-day off schedules ^16, 61–64^. However, the pharmacokinetic (PK) profile of CX-5461 in mice has yet to be reported in enough detail to judge the clinical relevance of such dosages and resultant plasma concentrations. To determine a PK-guided clinically relevant dosage, we measured plasma and tumor concentrations of CX-5461 in athymic nude mice at time points from 30 minutes to 24 hours after doses of 5 mg/kg IV and 25 mg/kg IP (Fig. S7A & S7B). We then compared CX-5461 plasma area under the concentration-time curve (AUC) and average plasma concentration (Cavg) values to those reported in the available human phase I clinical trial, which established a single agent maximum tolerated dose of 170 mg/m^2^ by intravenous infusion ^15^. We established a dosage of 5 mg/kg once daily by IP injection for 5 days, starting on day 1, with a second 5-day course starting on day 8, every 21 days. Our complete PK reports for all compounds screened in this study (including CX-5461) have been made freely available on the CSTN website (https://cstn.stjude.cloud/resources and https://www.stjude.org/content/dam/en_US/shared/www/research/cstn-appendix-standard-of-care-drug-regimens.pdf).

### Rescreen of GDSC cell lines with CX-5461

CX-5461 rescreening on a subset of GDSC cell lines was performed at the University of Chicago’s Cellular Screening Center using the CellTiter Glo assay. We rescreened the IMR-5, CHP-134, KELLY, BE(2)-M17, SK-N-SH, SK-N-FI, RS4-11, and H2452 cell lines. Cells were plated in a 96-well plate for one day then drug treated in triplicate for 72 hours at 2-fold serial dilutions at 10 concentrations from 5 µM to 0.000 µM. Luminescence was quantitated using a standard plate reader and IC_50_ values were estimated using a generalized logistic regression model implemented in the “drc” package in R.

### High-throughput drug combination (synergy) screening

CX-5461 in combination with each of AZD6738, AZD1390, and SN-38 were dispensed into 384-well flat-bottom tissue culture-treated solid white plates (Corning, Catalog# 3570) using a Labcyte Echo 555 acoustic Liquid Dispenser (Beckman Coulter Life Sciences, USA). Twelve-point dose-response curves were generated using 1:2 dilution intervals for each drug. The top concentrations of the drugs were derived from pre-determined IC_50_ values and were as follows: 311.4 nM SN-38, 1.89 µM CX-5461, 19.93 µM AZD6738, and 87.19 µM AZD1390. All plates contained two replicates each of the 144 pairwise combination matrices (all possible combinations of drug concentrations), the drugs alone without combination, DMSO-only (as a 0% cell death control), and the maximum concentration of each drug combined (as a 100% cell death control). The final DMSO concentration of all wells was 0.536%. Each plate was generated twice for a total of four replicates per drug combination per cell line. Into the compound-containing plates, the neuroblastoma cell lines CHP-134 (2000 cells/well), KELLY (2250 cells/well), BE(2)-M17 (1750 cells/well), and IMR-5 (2750 cells/well) were plated in volumes of 40 µL using a Multidrop Combi reagent dispenser (ThermoFisher Scientific, USA) and settled for 20 seconds at 100x g in an Eppendorf 5810 centrifuge. Plates were then incubated at 37°C in 5% CO_2_ for 96 hours in a Liconic STX220 incubator. Following incubation, plates were placed at room temperature for 20 minutes before viability assessment. 25 µL of CellTiter-Glo (Promega Catalog# G9241) was then added to each well to measure viability using a Multidrop Combi and the plates were incubated for an additional 25 minutes at room temperature. Luminescence was then detected with an EnVision 2102 Multilabel Plate Reader (PerkinElmer Life Sciences, USA).

### Drug synergy analysis

Raw luminescent data were imported into the R statistical environment version 4.0.2 (www.r-project.org). Background-subtracted values in raw luminescent units (RLU) were assigned to the appropriate drugs and concentrations, and each replicate was separated. All replicates were normalized to the mean of their respective inter-plate controls (DMSO for 0% cell death, and the maximum concentration of each drug combined for 100% cell death). Normalized drug-only data were fit with log-logistic regression to produce dose-response curves using the packages drc ^65^ and tidydrc (https://github.com/angelovangel/tidydrc). EC_50_ values for each drug’s curve were extrapolated and plotted. Matrices of the percent cell death values were constructed using means of normalized data from each of the four replicates per group as input. From these normalized values, synergy scores were calculated for all tested concentration combinations. Delta scores derived from the Zero Interaction Potency (ZIP) model ^45^ were used to score synergy across all samples using the SynergyFinder package ^66^. The most significant synergy (or antagonism) scores of drug pairs were directly compared to their corresponding single agents by referencing original normalized data of individual replicates and plotting mean values of percent cell death. From the resulting synergy matrices, both the highest-and lowest-scoring concentration pairs were extracted to represent the most significant synergy and antagonism, respectively. These scores were used to reference the expected percent cell death of a given combination (based on assumed non-interaction) to test whether the differences were statistically significant. *P*-values were calculated using a one sample t-test.

### TOP2A and TOP2B activity assays

Decatenation assay: 1 U of human topoisomerase II was incubated with 200 ng kDNA in a 30 μl reaction at 37 C for 30 minutes under the following conditions: 50 mM Tris HCl (pH 7.5), 125 mM NaCl, 10 mM MgCl_2_, 5 mM DTT, 0.5 mM EDTA, 0.1 mg/ml bovine serum albumin (BSA) and 1 mM ATP. Each reaction was stopped by the addition of 30 μl chloroform/iso-amyl alcohol (24:1) and 30 μl Stop Dye before being loaded on a 1.0% TAE gel run at 90V for 90 minutes. Bands were visualized by ethidium bromide staining for 15 minutes and destaining for 10 minutes. Gels were scanned using documentation equipment (GeneGenius, Syngene, Cambridge, UK) and decatenation levels were calculated from band data obtained using ImageJ. A second individual also quantitated these images independently using different gel scanning software (GeneTools, Syngene, Cambridge, UK) and obtained similar results. The activity of the enzymes was determined before the testing of the compound. The amount of topoisomerase II required for optimal decatenation was determined by titration. The controls and compounds were tested over a range of dilutions from 1.734 µM to 400 µM and added to the reaction before the addition of the enzyme. Final DMSO concentration in the assays was 10% (v/v). All experiments were performed five times and run on 1% agarose gels. Dose-response curves were estimated using a generalized logistic regression model fit to the data using the R package “drc”, from which IC_25_s and 95% confidence intervals were also calculated. Note that IC_25_s were calculated because a measurable IC_50_ was not achieved within the active drug concentration range for CX-5461 in the TOP2A assays; to fit the dose-response curves properly it was also necessary to filter those data, removing points beyond the activity range of CX-5461.

Relaxation Assays: 1 U of human topo II was incubated with 500 ng supercoiled pBR322 in a 30 μl reaction at 37 C for 30 minutes under the following conditions: 50 mM Tris HCl (pH 7.5), 125 mM NaCl, 10 mM MgCl2, 5 mM DTT, 0.5 mM EDTA, 0.1 mg/ml bovine serum albumin (BSA) and 1 mM ATP. Each reaction was stopped by the addition of 30 μl chloroform/iso-amyl alcohol (24:1) and 30 μl Stop Dye before being loaded on a 1.0% TAE gel run at 90V for 90 minutes. Bands were visualized by ethidium bromide staining for 15 minutes and destaining for 10 minutes. Gels were scanned using documentation equipment (GeneGenius, Syngene, Cambridge, UK) and relaxation levels were calculated from band data obtained with gel scanning software (GeneTools, Syngene, Cambridge, UK). The data for all topoisomerase activity assays were generated by Inspiralis (Innovation Centre, Norwich Research Park, Colney Lane, Norwich, UK) and the experimental details above were provided by Inspiralis. The data were analyzed by Inspiralis and reanalyzed by M.P and P.G, achieving consistent results.

### Ligand docking and molecular dynamics simulations

Published crystal structures of etoposide-bound TOP2A (PDB: 5GWK) or TOP2B (PDB: 3QX3) were loaded into Maestro software (Schrödinger Release 2020-3). To prepare the protein for docking and simulations, the protein preparation wizard was used to assign bond orders, add hydrogens, create zero-order bonds to metals, create disulfide bonds, and fill in missing side chains and loops. Default parameters were used for the optimization of hydrogen-bond assignment (sampling of water orientations and use of pH 7.0). Waters beyond 5 Å of het groups or with fewer than three hydrogen bonds to non-waters were removed. Restrained energy minimization was applied using the OPLS3e forcefield. Prepared protein systems were further checked by Ramachandran plots, ensuring there were no steric clashes.

For docking CX-5461 into TOP2A and TOP2B, the 3D structure of CX-5461 was first obtained from the PubChem database (www.pubchem.ncbi.nlm.nih.gov). The flexible ligand alignment tool in Maestro was used to align the core scaffold of CX-5461 to etoposide based on common scaffolds defined by the Bermis-Murcko method. The aligned molecule was then translated into the etoposide bound site based on these aligned scaffolds. The ligand position was adjusted in the binding site so that CX-5461 would occupy the optimal volume of the site through rigid body minimization of CX-5461 using Prime. The Prime tool was further used to adjust protein sidechain rotamers of TOP2A and TOP2B that are within 5 Å of CX-5461. The binding site was further optimized by administering a conjugate-gradient minimization algorithm for all residues within 5 Å of CX-5461, with a 200kcal/mol/Å^2^ restraint placed on a shell of residues greater than 5 Å but less than 7 Å from CX-5461. All atoms greater than 7 Å from CX-5461 were ignored by the algorithm.

For molecular dynamics simulations, systems were built for CX-5461-bound TOP2A and TOP2B by using the system builder panel of Desmond (Schrödinger Release 2020-3). The SPC solvent model was used, and the forcefield was set to OPLS3e. Solvated systems were loaded into the workspace by using the molecular dynamics panel. The total simulation time for each system was set to 10 nanoseconds, with 100-picosecond trajectory recording intervals. The system energy was set to 1.2, and the ensemble class used was NPT. Simulations were set to run at 300.0 K and 1.01325 bar. The option to relax model systems before simulations was selected.

Simulations were clustered based on RMSD using default parameters in the trj_cluster.py command line script available in Schrodinger utilities. This resulted in 12 clusters for TOP2A and 16 clusters for TOP2B. For each cluster, the representative structure file was used to calculate the relative binding affinity of TOP2A and TOP2B to the CX-5461 ligand. The binding energy was calculated using the Prime MM-GBSA tool from the Maestro GUI. The population of each cluster was used to determine weights for calculating a weighted average binding energy for TOP2A and TOP2B with CX-5461 from each of the clusters of the trajectory.

### Cell line drug screening analysis

Preprocessed GDSC data (drug sensitivity, gene expression, and mutation calls) were obtained from the GDSC website (www.cancerrxgene.org). All analyses of these data were performed using R and figures were created using the base graphics functions and the ggplot2 package. Basic statistical tests such as t-tests, linear regression, Wilcoxon-rank sum tests, and correlation tests were performed using the base statistical functions in R and, where appropriate, *P*-values were corrected for multiple testing by estimating false-discovery rates using the Benjamini and Hochberg method. The PRISM drug screening data were obtained from www.depmap.org.

### Patient *-omics* data analysis

Neuroblastoma patient tumor data was obtained from ^41^ who aggregated data from multiple previously published neuroblastoma studies including RNA-seq, clinical annotations, and genomics sequencing data. The associations between *TOP2B* expression and individual genomic and clinical features were estimated using linear regression. Sparse regression models, aimed at identifying independent predictors of *TOP2B* expression were fit using ElasticNet regression, implemented in the R package *glmnet*, where the alpha parameter was fixed to 0.5 and the lambda parameter was deduced by 10-fold cross-validation. GTEx data were obtained from the GTEx data portal (www.gtexportal.org), TCGA data from the Genomics Data Commons, paired relapse data was obtained from TARGET ^67^ and data from paired primary tumor and metastatic sites was obtained from ^42^. Paired patient tumor – PDX gene expression data for our 2 xenografts was obtained from the St. Jude Cloud ^68^. When comparing the expression of *TOP2B*, we also normalized these gene expression data by calculating their log_2_ fold-change relative to housekeeping genes *ACTIN* and *GAPDH*, which are treated as negative controls whose expression is not expected to vary, thus mitigating the effect of gene expression differences that would be expected to arise due to dataset-specific effects.

### Computational alignment of cell lines and tumor samples

To align cell lines to their corresponding lineages of patient tumor samples, the Celligner tool ^39^ was used. Clinical expression data were obtained from the UCSC Xena data hub (www.xena.ucsc.edu), which included processed data for tumor samples originating from The Cancer Genome Atlas (TCGA, www.cancer.gov) and TARGET (www.ocg.cancer.gov) datasets. Cell line expression data were obtained from Expression Atlas (www.ebi.ac.uk), which included RNA-sequencing of cell lines used in the Sanger Genomics of Drug Sensitivity in Cancer (GDSC, www.cancerrxgene.org) project. All expression data were in units of log_2_(TPM+1) and datasets were filtered such that only common lineages between tumors and cell lines were kept and used as input. After completion of the Celligner tool and the resulting UMAP plot was generated, alignments of cell lines to tumors were quantified by their respective lineages. Briefly, Celligner-generated tumor clusters were isolated and kept if they represented ≥ 80% of a given lineage. Proportions of cell lines that correctly aligned to their tumor counterpart clusters (by lineage) were then calculated. Fisher exact tests were performed to assess statistical significance for all alignments, and the resulting *P*-values and odds ratios were generated for each lineage. Step-by-step details of the Celligner tool, post-alignment analysis, and all input data are available in the Open Science Framework (www.osf.io/pzke4).

### Materials availability

PDX models that were used in this study are freely available upon request from the Childhood Solid Tumor Network (CSTN), which is hosted at St. Jude (https://www.stjude.org/research/resources-data/childhood-solid-tumor-network/available-resources.html). We have also made the full detailed pharmacokinetic reports for the compounds used in our PDX studies (including CX-5461) available on the CSTN website (https://cstn.stjude.cloud/resources and https://www.stjude.org/content/dam/en_US/shared/www/research/cstn-appendix-standard-of-care-drug-regimens.pdf).

### Data and code availability

The code to reproduce computational analyses in this study, including the supporting data, have been deposited on Open Science Framework (osf.io/pzke4/).

