## Supplementary Figures for "The chemotherapeutic CX-5461 primarily targets TOP2B and exhibits selective activity in high-risk neuroblastoma"

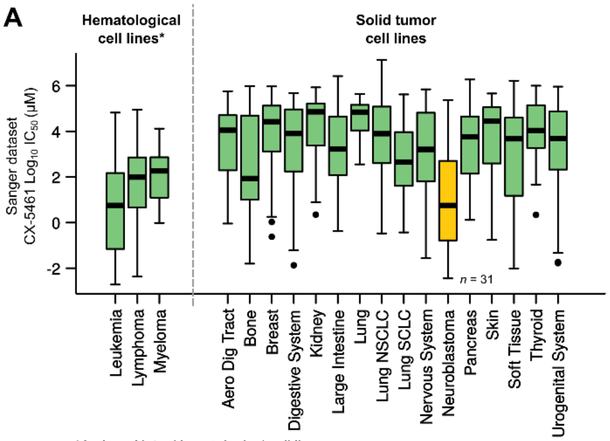


*Authors Note: Hematological cell lines are generally more drug sensitive in this dataset, thus the low $IC_{50}$ values in hematological cell lines do not imply selective activity.

**B**

| Cell line | IC50 (µM) CX-5461 | P53 | MYCN Copy Number |
|---|---|---|---|
| IMR-5 | 0.018 | WT | 14 |
| CHP-134 | 0.041 | WT | 12 |
| KELLY | 0.589 | Mut | 14 |
| BE(2)-M17 | 5.491 | Mut | 14 |
| SK-N-SH | 0.220 | Mut | 3 |
| SK-N-FI | 27.267 | Mut | 2 |
| RS4-11 | 0.0684 | Sensitive cell control | |
| H2452 | 31.5601 | Resistant cell control | |

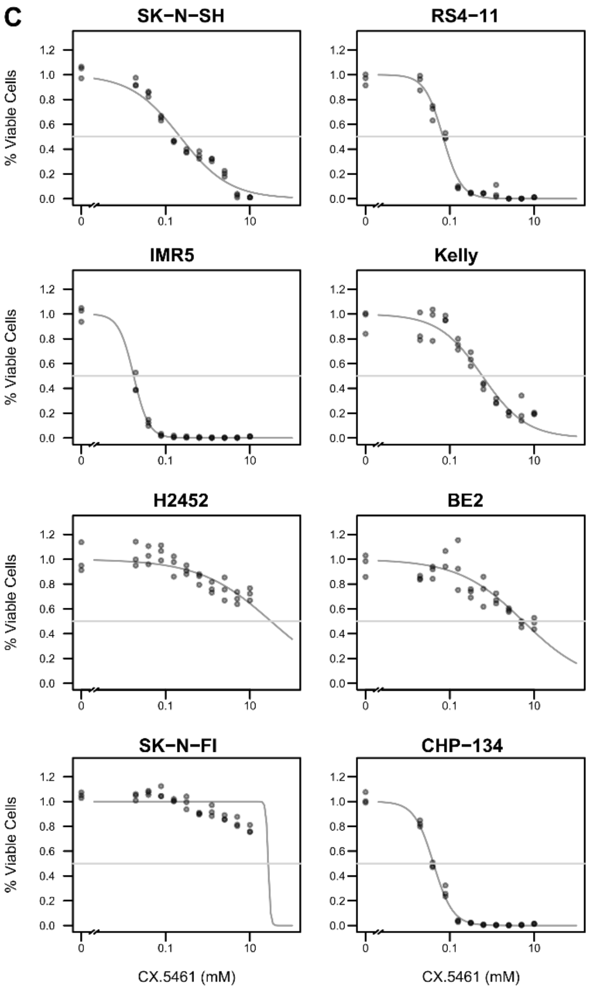


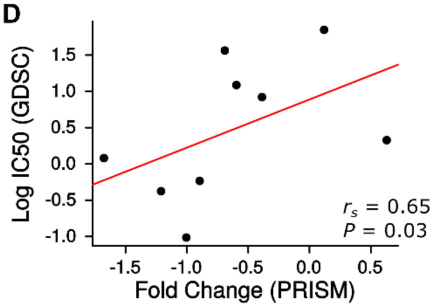


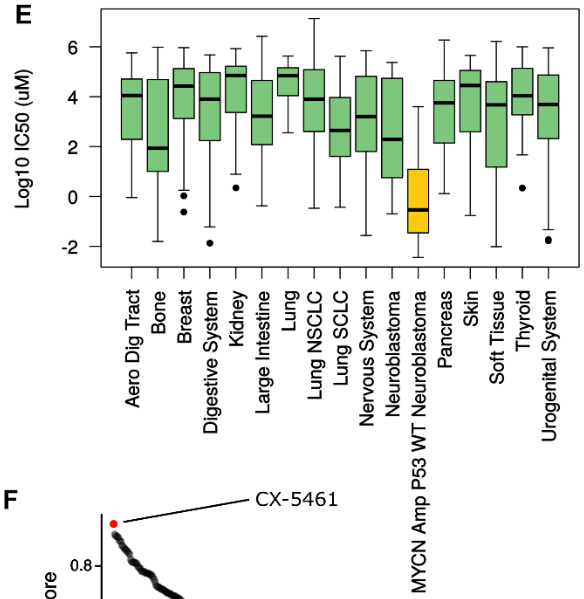


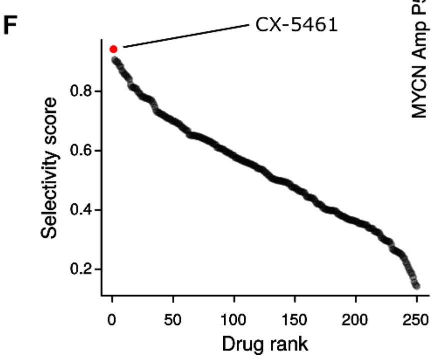

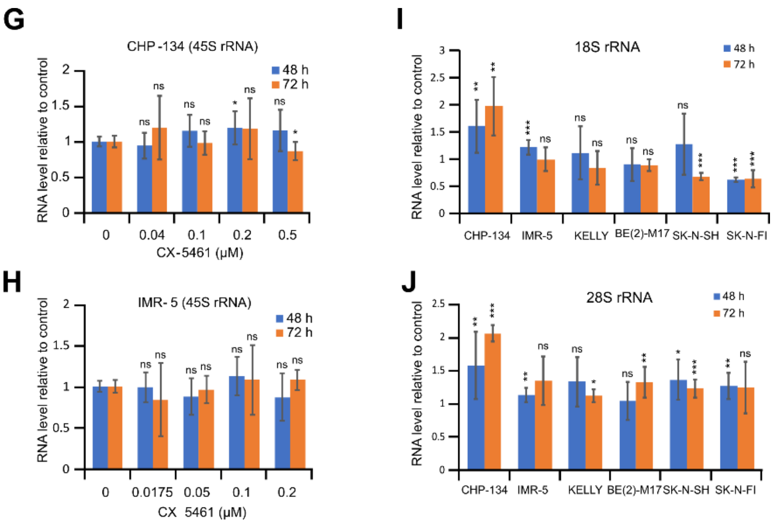


**Figure S1**

A. Boxplot of all $IC_{50}$ values for CX-5461 from the 2016 release of the GDSC dataset with hematological cancer cell lines (Iorio et al, Cell 2016). Note that hematological cell lines tend to be generally more drug sensitive in these screens and a low $IC_{50}$ compared to other cell lines does not imply selective killing.

B. Table showing $IC_{50}$ values obtained and other relevant details from our re-screen of a subset of 8 GDSC cell lines with CX-5461.

C. Cell viability data for the 8 cell lines, showing the percentage of viable cells (y-axis) against the concentration of CX-5461 (x-axis). Dose-response curves were estimated using a generalized logistic regression model fit to the data using the R package “drc”.

D. Scatterplot of log $IC_{50}$ values from GDSC (y-axis) plotted against fold change values in PRISM (x-axis) for CX-5461 for the 9 neuroblastoma cell lines that were screened in both datasets; the *P*-value was estimated using a one-sided Spearman's Rank Correlation test.

E. Boxplot of $\log_{10}(IC_{50})$ values for CX-5461 from all solid tumor cell lines in GDSC. Hematological cancers are not included. *MYCN* amplified *TP53* WT neuroblastoma cell lines have been separated from other neuroblastoma cell lines.

F. Waterfall plot showing Selectivity Scores (y-axis) for *MYCN* amplified *TP53* WT neuroblastoma cell lines for 265 compounds screened by GDSC.

G & H. 45S pre-rRNA expression levels (y-axis) in CX-5461 treated CHP-134 and IMR-5 cells were measured by qPCR. Data represent mean ± SD of 3 independent experiments. *$P < 0.05$; ns, no significant difference.

I & J. 18S and 28S rRNA expression (y-axis) in CX-5461 treated neuroblastoma cell lines. Data represent mean ± SD of 3 independent experiments. *$P < 0.05$; **$P < 0.01$; ***$P < 0.001$; ns, no significant difference. CX-5461 concentration: CHP-134, 0.2 µM; IMR-5, 0.05 µM; KELLY, 2 µM; BE(2)-M17, 10 µM; SK-NSH, 2 µM; SK-N-FI, 20 µM.

A

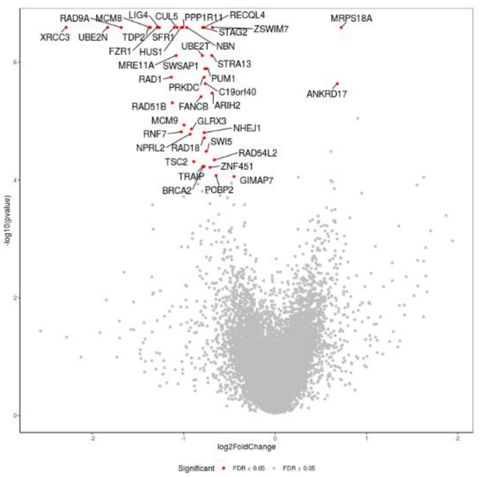


B

| Index | Name | P-value | Adjusted p-value | Odds Ratio | Combined score |
|---|---|---|---|---|---|
| 1 | recombinational repair (GO:0000725) | 1.067e-18 | 2.723e-15 | 77.46 | 3205.61 |
| 2 | double-strand break repair via homologous recombination (GO:0000724) | 2.371e-18 | 4.033e-15 | 72.37 | 2936.94 |
| 3 | DNA recombinase assembly (GO:0000730) | 4.121e-7 | 0.0002337 | 187.50 | 2756.62 |
| 4 | DNA repair complex assembly (GO:0090735) | 4.121e-7 | 0.0002103 | 187.50 | 2756.62 |
| 5 | double-strand break repair via synthesis-dependent strand annealing (GO:0045003) | 4.121e-7 | 0.0001912 | 187.50 | 2756.62 |
| 6 | double-strand break repair (GO:0006302) | 8.503e-23 | 4.339e-19 | 53.19 | 2703.14 |
| 7 | strand invasion (GO:0042148) | 0.00008138 | 0.01978 | 142.86 | 1345.19 |
| 8 | response to ionizing radiation (GO:0010212) | 1.597e-10 | 1.164e-7 | 45.45 | 1025.36 |
| 9 | telomeric loop disassembly (GO:0090657) | 0.0001737 | 0.03410 | 100.00 | 865.80 |
| 10 | DNA recombination (GO:0006310) | 7.983e-8 | 0.00005092 | 45.45 | 742.88 |

C

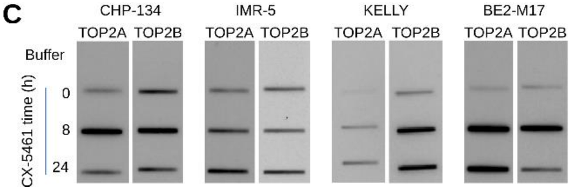


**Figure S2.**

A. Volcano plot showing the results of our genome wide CRISPR knockout screen in CHP-134 cells. The x-axis shows the mean fold change for the normalized abundance of the guide RNAs targeting each gene in CX-5461 compared to DMSO treated control cells. P-values are shown on the y-axis and significant knockouts, at a false discovery rate of <5%, are highlighted in red.
B. Table of the top pathways from our genome wide CRISPR knockout screen in CHP-134 cells as reported by the Enrichr software for genes at an FDR <5%.
C. RADAR assay in neuroblastoma cells. Cells were treated with CX-5461 or DMSO (0 h) for the time indicated. DNA – topoisomerase covalent complexes were extracted and visualized using TOP2A and TOP2B antibodies. CX-5461 concentration used for each cell line: CHP-134, 0.2 µM; IMR-5, 0.05 µM; KELLY, 2 µM; BE(2)-M17, 10 µM.

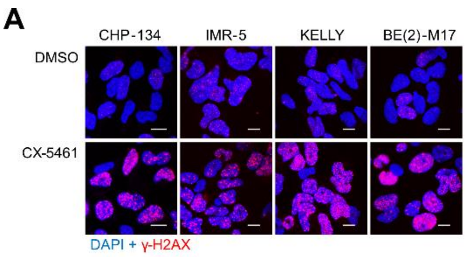


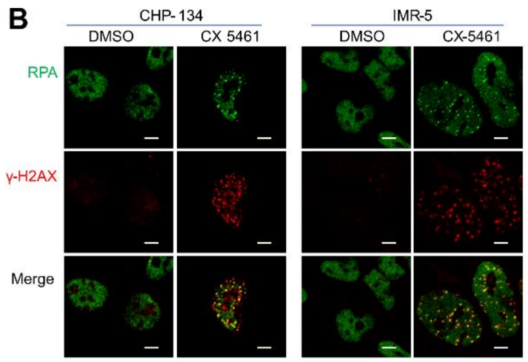


C

| Time | 0 | 30 min | | 24 h | |
|---|---|---|---|---|---|
| | Mean | Mean | *p* | Mean | *p* |
| Tail length | 7.30 | 18.87 | 5.24E-16 | 26.75 | 1.27E-25 |
| Tail DNA percent | 7.53 | 20.78 | 2.94E-10 | 28.13 | 2.82E-15 |
| Tail moment | 1.00 | 7.35 | 6.42E-09 | 12.86 | 2.27E-16 |

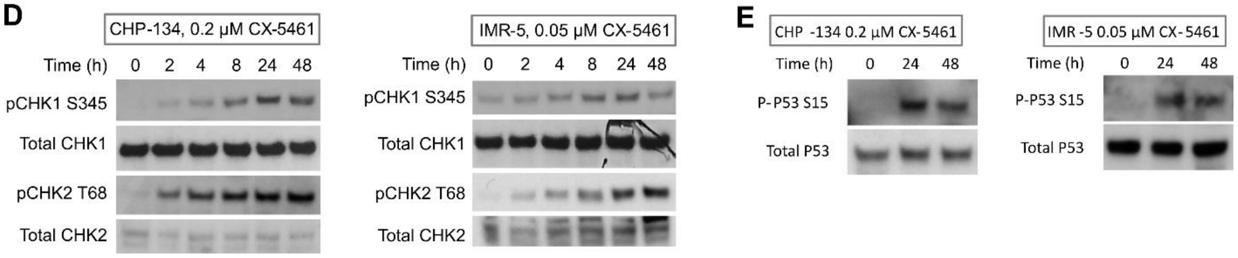


**Figure S3.**

A. CX-5461 induced γ-H2AX foci in neuroblastoma cell lines. Cells were treated with DMSO or CX-5461 for 24 h. Nuclei were stained with DAPI (blue) and phosphorylated histone H2AX was stained with γ-H2AX antibody (red). CX-5461 concentration: CHP-134, 0.2 μM; IMR-5, 0.05 μM; KELLY, 2 μM; BE(2)-M17, 0.5 μM. Scale bar = 10 μm.

B. CX-5461 induced RPA and γ-H2AX foci. After treated with DMSO or CX-5461 (0.2 μM for CHP-134 and 0.05 μM for IMR-5) for 24 h, cells were stained with RPA (green) and γ-H2AX (red) antibodies for immunofluorescence. Scale bar = 5 μm.

C. Quantification of comet assay in Figure 3F. Tail length, tail DNA percent, and tail moment were calculated with OpenComet image analysis tool. *P*-values are from 2-tailed t-test.

D. Western blots for CHK1, CHK2 and phosphorylated CHK1 and CHK2. Cells were treated with CX-5461 for the time and concentration indicated.

E. Western blots for p53 and phosphorylated p53. Cells were treated with CX-5461 for the time and concentration indicated.

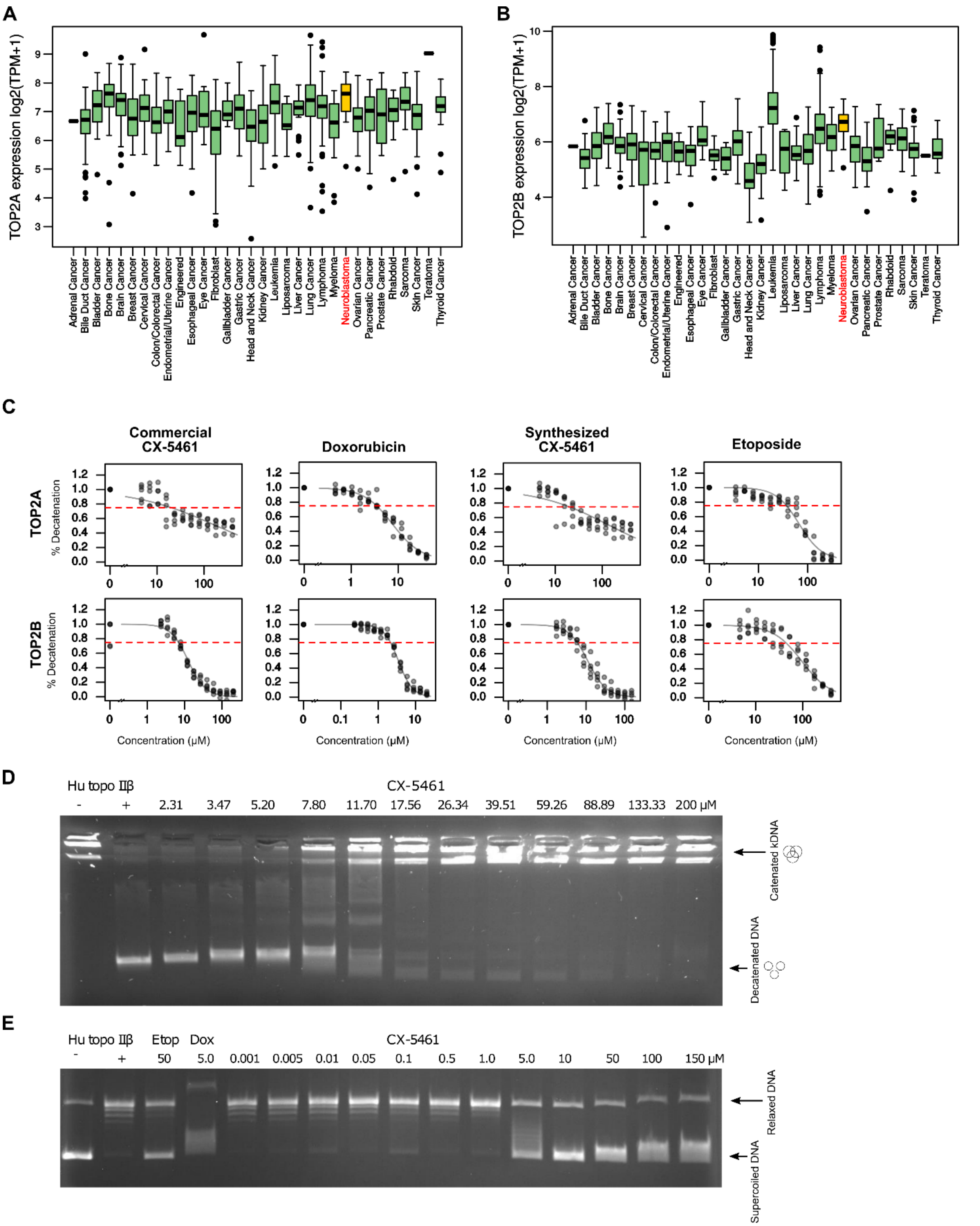


**Figure S4**

A. Boxplot showing the gene expression level (y-axis) of TOP2A in all DepMap cancer cell lines, 20Q2 release of these RNA-seq data. Data obtained from www.depmap.org.

B. Boxplot showing the gene expression level (y-axis) of TOP2B in all DepMap cancer cell lines, 20Q2 release of these RNA-seq data. Data obtained from www.depmap.org.
C. Summarized data from all decatenation assays for the four drugs indicated, for both TOP2A (top row) and TOP2B (bottom row) assays, showing the percentage of decatenated kDNA (y-axes) against the drug concentration (x-axes). Dose-response curves were estimated using a generalized logistic regression model fit to the data using the R package “drc”, from which $IC_{25}$s and 95% confidence intervals were also calculated. Note that $IC_{25}$s were calculated because a reliable $IC_{50}$ was not achieved within the active drug concentration range for CX-5461 in the TOP2A assays.
D. Representative gel image from a single run of the TOP2B decatenation assay for CX-5461. 1 U of purified human TOP2B protein was incubated with 200 ng of kinetoplast DNA (kDNA) with the indicated drug concentration for 30 minutes at 37C. The product of this reaction was run on a 1% TAE gel (details in Methods). Gels were scanned (GeneGenius, Syngene, UK) and quantified using ImageJ.
E. Representative gel image from a single run of the TOP2B relaxation assay for CX-5461.

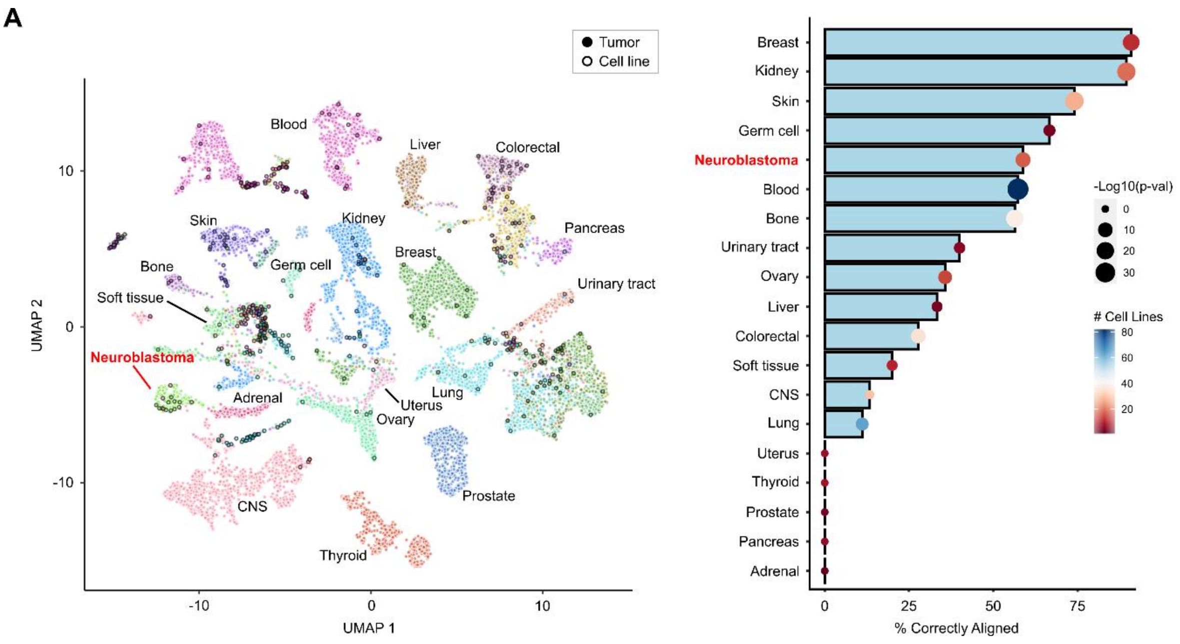
A
Tumor
Cell line
Blood
Liver
Colorectal
Skin
Kidney
Pancreas
Bone
Germ cell
Breast
Urinary tract
Soft tissue
Neuroblastoma
Adrenal
Uterus
Lung
Ovary
Prostate
CNS
Thyroid
UMAP 1
UMAP 2
10
0
-10
Breast
Kidney
Skin
Germ cell
Neuroblastoma
Blood
Bone
Urinary tract
Ovary
Liver
Colorectal
Soft tissue
CNS
Lung
Uterus
Thyroid
Prostate
Pancreas
Adrenal
0
25
50
75
% Correctly Aligned
-Log10(p-val)
0
10
20
30
### Cell Lines
80
60
40
20

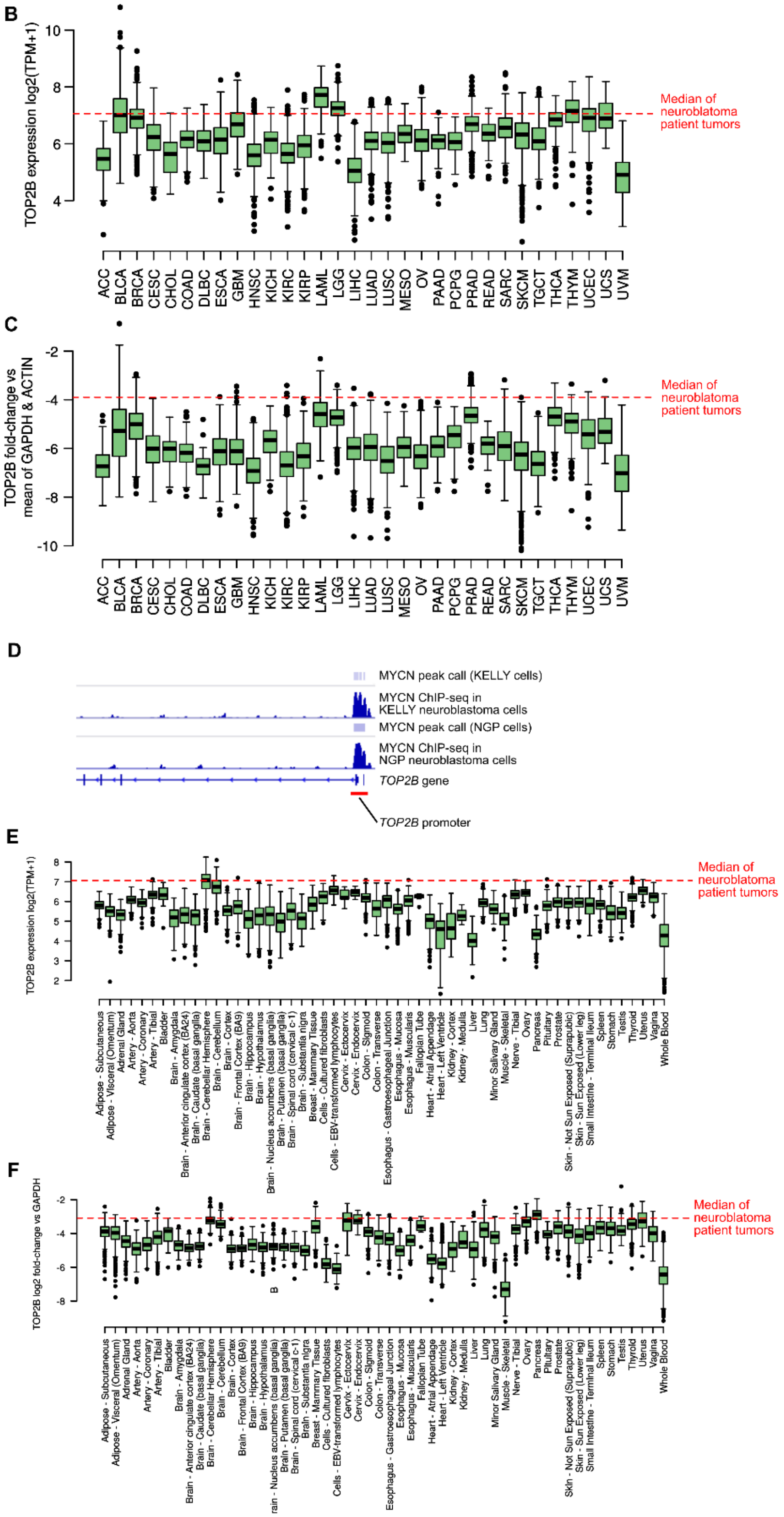
B
TOP2B expression log2(TPM+1)
10
8
6
4
Median of neuroblatoma patient tumors
ACC BLCA BRCA CESC CHOL COAD DLBC ESCA GBM HNSC KICH KIRC KIRP LAML LGG LIHC LUAD LUSC MESO OV PAAD PCPG PRAD READ SARC SKCM TGCT THCA THYM UCEC UCS UVM
C
TOP2B fold-change vs mean of GAPDH & ACTIN
-2
-4
-6
-8
-10
Median of neuroblatoma patient tumors
ACC BLCA BRCA CESC CHOL COAD DLBC ESCA GBM HNSC KICH KIRC KIRP LAML LGG LIHC LUAD LUSC MESO OV PAAD PCPG PRAD READ SARC SKCM TGCT THCA THYM UCEC UCS UVM
D
MYCN peak call (KELLY cells)
MYCN ChIP-seq in KELLY neuroblastoma cells
MYCN peak call (NGP cells)
MYCN ChIP-seq in NGP neuroblastoma cells
TOP2B gene
TOP2B promoter
E
TOP2B expression log2(TPM+1)
8
7
6
5
4
3
2
Median of neuroblatoma patient tumors
Adipose - Subcutaneous
Adipose - Visceral (Omentum)
Adrenal Gland
Artery - Aorta
Artery - Coronary
Artery - Tibial
Bladder
Brain - Amygdala
Brain - Anterior cingulate cortex (BA24)
Brain - Caudate (basal ganglia)
Brain - Cerebellar Hemisphere
Brain - Cerebellum
Brain - Cortex
Brain - Frontal Cortex (BA9)
Brain - Hippocampus
Brain - Hypothalamus
Brain - Nucleus accumbens (basal ganglia)
Brain - Putamen (basal ganglia)
Brain - Spinal cord (cervical c-1)
Brain - Substantia nigra
Breast - Mammary Tissue
Cells - Cultured fibroblasts
Cells - EBV-transformed lymphocytes
Cervix - Ectocervix
Cervix - Endocervix
Colon - Sigmoid
Colon - Transverse
Esophagus - Gastroesophageal Junction
Esophagus - Mucosa
Esophagus - Muscularis
Fallopian Tube
Heart - Atrial Appendage
Heart - Left Ventricle
Kidney - Cortex
Kidney - Medulla
Liver
Lung
Minor Salivary Gland
Muscle - Skeletal
Nerve - Tibial
Ovary
Pancreas
Pituitary
Prostate
Skin - Not Sun Exposed (Suprapubic)
Skin - Sun Exposed (Lower leg)
Small Intestine - Terminal Ileum
Spleen
Stomach
Testis
Thyroid
Uterus
Vagina
Whole Blood
F
TOP2B log2 fold-change vs GAPDH
-2
-4
-6
-8
Median of neuroblatoma patient tumors
Adipose - Subcutaneous
Adipose - Visceral (Omentum)
Adrenal Gland
Artery - Aorta
Artery - Coronary
Artery - Tibial
Bladder
Brain - Amygdala
Brain - Anterior cingulate cortex (BA24)
Brain - Caudate (basal ganglia)
Brain - Cerebellar Hemisphere
Brain - Cerebellum
Brain - Cortex
Brain - Frontal Cortex (BA9)
Brain - Hippocampus
Brain - Hypothalamus
rain - Nucleus accumbens (basal ganglia)
Brain - Putamen (basal ganglia)
Brain - Spinal cord (cervical c-1)
Brain - Substantia nigra
Breast - Mammary Tissue
Cells - Cultured fibroblasts
Cells - EBV-transformed lymphocytes
Cervix - Ectocervix
Cervix - Endocervix
Colon - Sigmoid
Colon - Transverse
Esophagus - Gastroesophageal Junction
Esophagus - Mucosa
Esophagus - Muscularis
Fallopian Tube
Heart - Atrial Appendage
Heart - Left Ventricle
Kidney - Cortex
Kidney - Medulla
Liver
Lung
Minor Salivary Gland
Muscle - Skeletal
Nerve - Tibial
Ovary
Pancreas
Pituitary
Prostate
Skin - Not Sun Exposed (Suprapubic)
Skin - Sun Exposed (Lower leg)
Small Intestine - Terminal Ileum
Spleen
Stomach
Testis
Thyroid
Uterus
Vagina
Whole Blood

**Figure S5**

A. UMAP representation showing the genome-wide gene-expression-based alignment of all GDSC (Sanger) cell lines with patient tumor gene expression data from TCGA, Treehouse and TARGET, generated using the Celligner tool. The points representing individual cell lines have a black border. Points have been colored by their lineage and clusters have been labelled by tumor lineage. The bar plot (right panel) shows the proportion of cell lines from a given lineage that correctly align with the appropriate patient tumor cluster (x-axis). The points have been colored by the number of cell lines in the datasets. The size of the points have been scaled by the *P*-value obtained from a Fisher's exact test for cluster membership.
B. The median *TOP2B* $\log_2$(TPM+1) normalized expression from 88 neuroblastoma patient tumors compared to all TCGA lineages.
C. The same data as (A) but with *TOP2B* normalized to *ACTIN* and *GAPDH* housekeeping genes.
D. ChIP-seq data for MYCN at the TOP2B locus and promoter. Data shown from NGP and KELLY neuroblastoma cell lines. This figure was created from published ChIP-seq data generated by Zeid *et al.* (Nature Genetics 2018).
E. The median *TOP2B* $\log_2$(TPM+1) normalized expression from 88 neuroblastoma patient tumors compared to all GTEx tissue types.
F. The same data as (D) but with *TOP2B* normalized to *GAPDH* housekeeping genes (note that *ACTIN* expression was not used as a negative control here as it has variable expression across GTEx normal tissues).

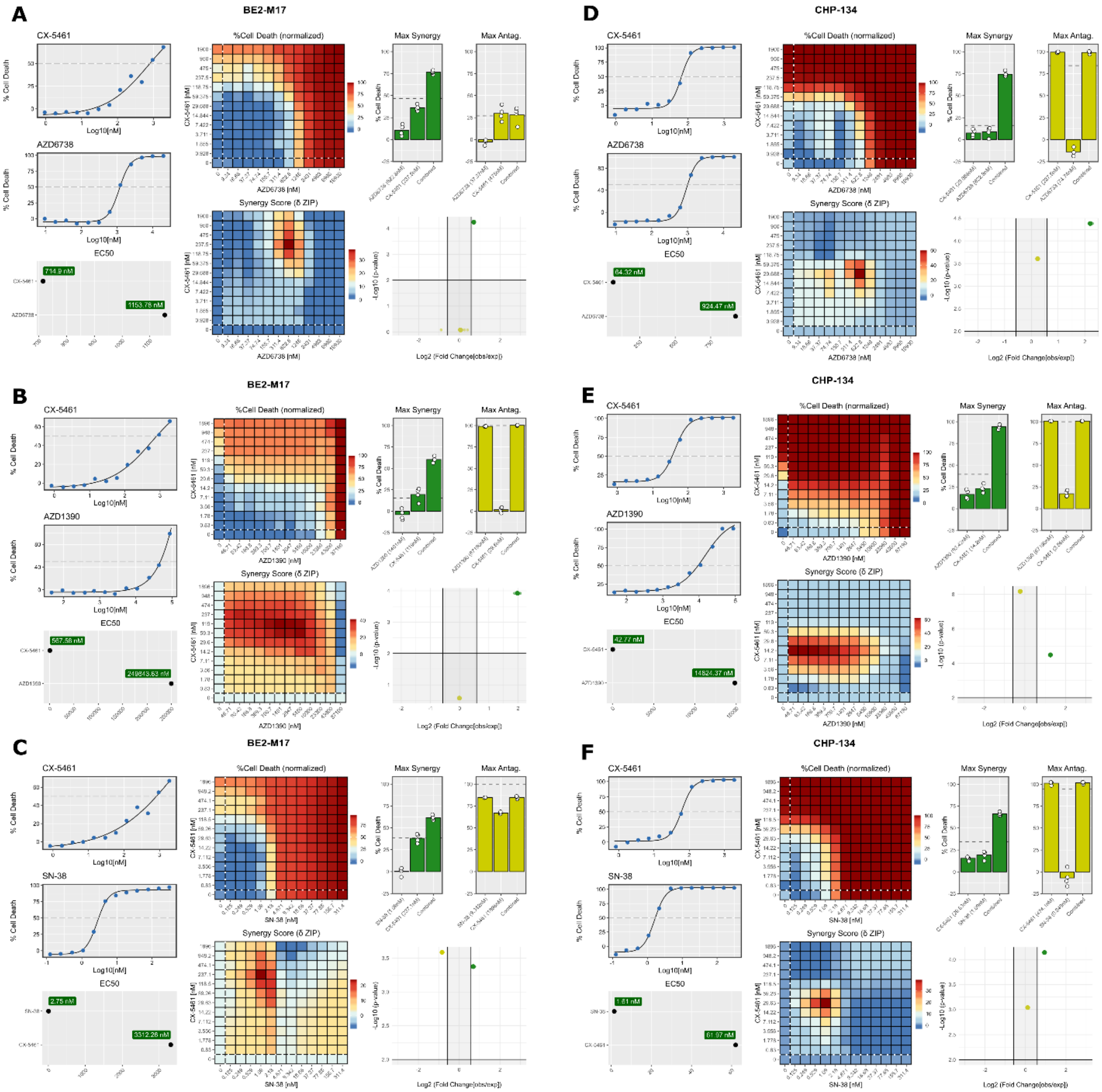
A
BE2-M17
CX-5461
AZD6738
EC50
714.9 nM
1153.76 nM
%Cell Death (normalized)
Synergy Score (δ ZIP)
Max Synergy
Max Antag.
% Cell Death
Log10[nM]
AZD6738 [nM]
CX-5461 [nM]
-Log10 (p-value)
Log2 (Fold Change[obs/exp])
B
BE2-M17
CX-5461
AZD1390
EC50
567.56 nM
249843.63 nM
AZD1390 [nM]
C
BE2-M17
CX-5461
SN-38
EC50
2.75 nM
3312.26 nM
SN-38 [nM]
D
CHP-134
CX-5461
AZD6738
EC50
64.32 nM
924.47 nM
E
CHP-134
CX-5461
AZD1390
EC50
42.77 nM
14824.37 nM
F
CHP-134
CX-5461
SN-38
EC50
1.81 nM
61.97 nM

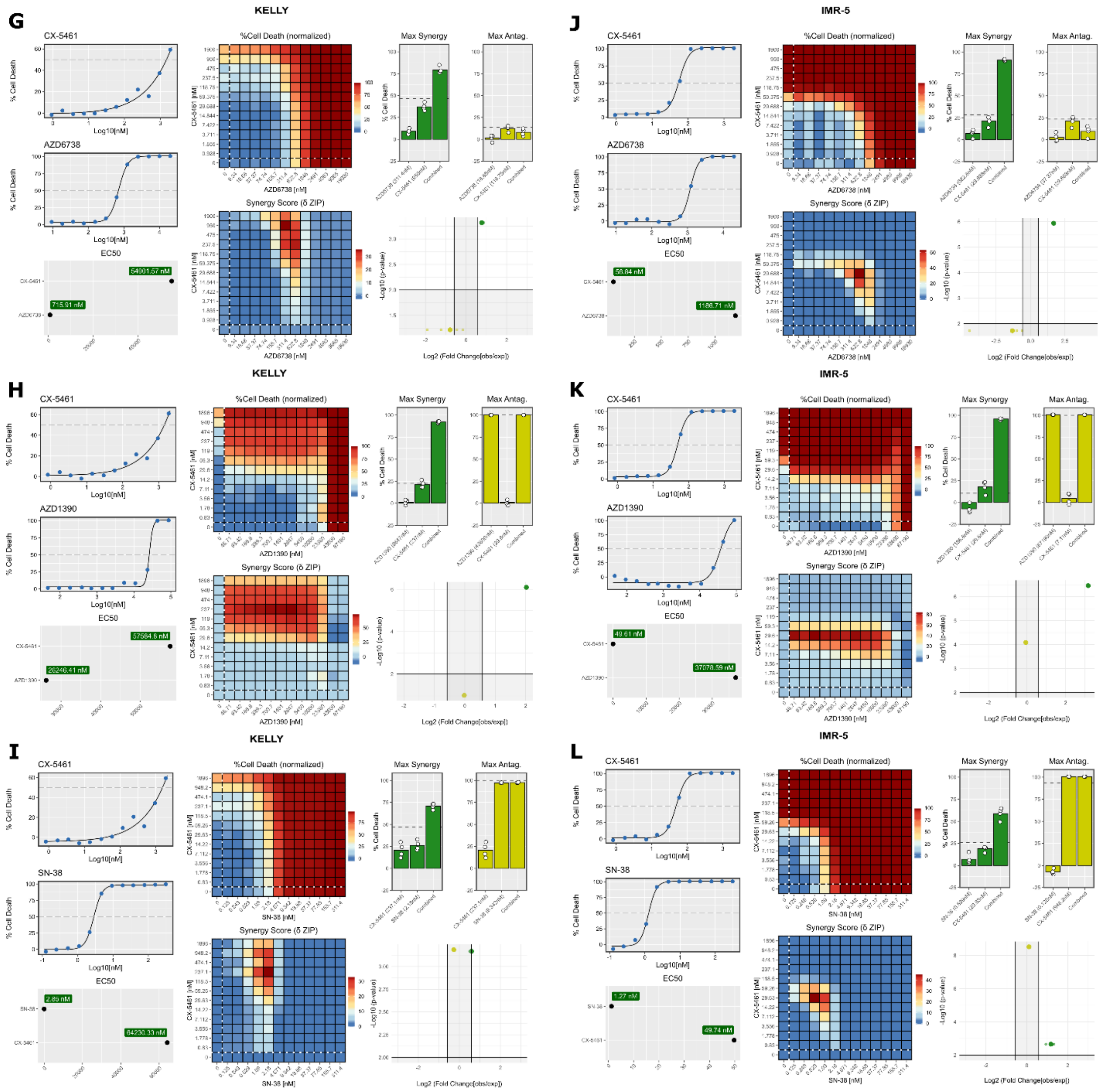


#### Figure S6

A-C. Drug screening with combination of CX-5461 and AZD-6738 (A), AZD-1390 (B), or SN-38 (C) in BE(2)-M17 cells. Dose-response curves and extrapolated $EC_{50}$ are shown on the left. Heatmap matrices (middle) represent percent cell death (upper) and calculated synergy scores (lower) of the combination treatment. Bar plots (upper right) represent the maximum synergy (green) and maximum antagonism (yellow) scores corresponding to the synergy matrix. White dots represent four independent experiments corresponding to these maxima. Volcano plot (lower right) represents the $\log_2$ fold change (x-axis) of the observed vs. expected (based on an additive model) values and *P*-values derived from one-sample t-tests (y-axis). Larger opaque dots represent the average of four independent experiments, while the smaller translucent dots correspond to the individual replicates.

D-F. Same as (A), (B), (C) respectively but in CHP-134 cells.
G-I. Same as (A), (B), (C) respectively but in KELLY cells.
J-L. Same as (A), (B), (C) respectively but in IMR-5 cells.

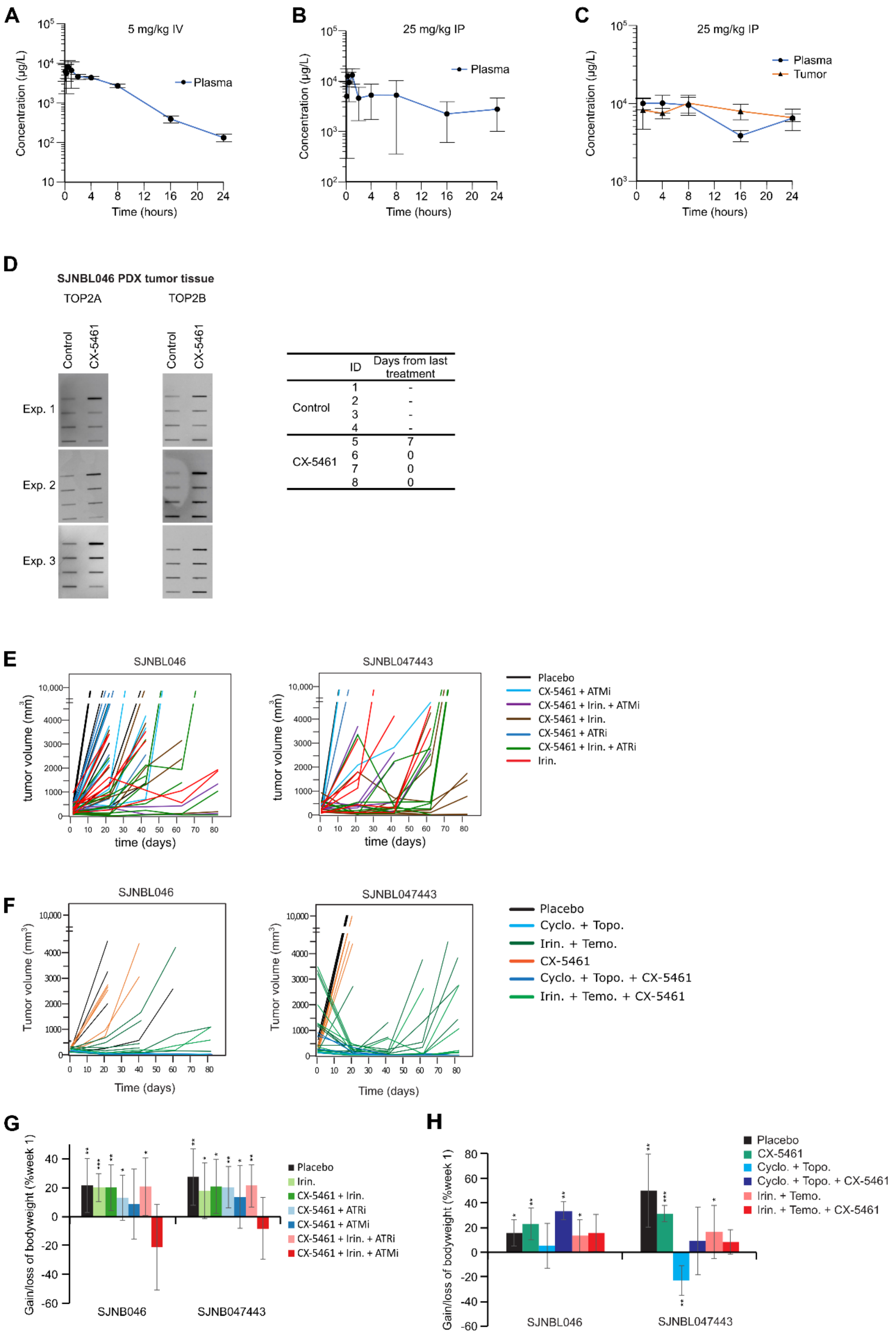
A
5 mg/kg IV
Plasma
Concentration (µg/L)
Time (hours)
B
25 mg/kg IP
Plasma
Concentration (µg/L)
Time (hours)
C
25 mg/kg IP
Plasma
Tumor
Concentration (µg/L)
Time (hours)
D
SJNBL046 PDX tumor tissue
TOP2A
TOP2B
Control
CX-5461
Exp. 1
Exp. 2
Exp. 3
ID
Days from last treatment
Control 1 -
2 -
3 -
4 -
CX-5461 5 7
6 0
7 0
8 0
E
SJNBL046
SJNBL047443
tumor volume (mm3)
time (days)
Placebo
CX-5461 + ATMi
CX-5461 + Irin. + ATMi
CX-5461 + Irin.
CX-5461 + ATRi
CX-5461 + Irin. + ATRi
Irin.
F
SJNBL046
SJNBL047443
Tumor volume (mm3)
Time (days)
Placebo
Cyclo. + Topo.
Irin. + Temo.
CX-5461
Cyclo. + Topo. + CX-5461
Irin. + Temo. + CX-5461
G
Gain/loss of bodyweight (%week 1)
SJNB046
SJNB047443
Placebo
Irin.
CX-5461 + Irin.
CX-5461 + ATRi
CX-5461 + ATMi
CX-5461 + Irin. + ATRi
CX-5461 + Irin. + ATMi
H
Gain/loss of bodyweight (%week 1)
SJNBL046
SJNBL047443
Placebo
CX-5461
Cyclo. + Topo.
Cyclo. + Topo. + CX-5461
Irin. + Temo.
Irin. + Temo. + CX-5461

**Figure S7**

A. Plasma concentration (y-axis) of CX-5461 achieved in athymic nude mice over 24 hours (x-axis) following a 5 mg/kg intravenous (IV) injection of CX-5461 (n = 3 mice at each time point). Data represent mean ± SD.
B. Same as (A) but mice were treated with 25 mg/kg CX-5461 via intraperitoneal (IP) injection.
C. Plasma and tumor concentrations (y-axis) of CX-5461 achieved in athymic nude mice over 24 hours (x-axis) following a 25 mg/kg IP injection of CX-5461 (n = 3 mice at each time point). Data represent mean ± SD.
D. Raw images for data in Figure 7A. The table (right) shows the duration between the harvesting of each tumor and the previous drug treatment for each mouse.
E. Tumor volume (y-axis) measured by ultrasound in SJNBL046 (left) and SJNBL047443 (right) PDX mice in study 1. Curves are colored by treatment group.
F. Tumor volume (y-axis) measured by ultrasound in SJNBL046 (left) and SJNBL047443 (right) PDX mice in study 2. Curves are colored by treatment group.

G-H. Bar plot showing % change of bodyweight at death for all mice in each group (after completion of therapy, body weight of all mice survived were monitored till death) in study 1 (G) and study 2 (H). Data represent the mean ± SD. *$P$ < 0.05, **$P$ < 0.01, ***$P$ < 0.001.
